# Krause corpuscles of the genitalia are vibrotactile sensors required for normal sexual behavior

**DOI:** 10.1101/2023.06.14.545006

**Authors:** Lijun Qi, Michael Iskols, Annie Handler, David D. Ginty

**Affiliations:** Department of Neurobiology, Howard Hughes Medical Institute, Harvard Medical School, 220 Longwood Avenue, Boston, MA 02115

## Abstract

Krause corpuscles, first discovered in the 1850s, are enigmatic sensory structures with unknown physiological properties and functions found within the genitalia and other mucocutaneous tissues. Here, we identified two distinct somatosensory neuron subtypes that innervate Krause corpuscles of the mouse penis and clitoris and project to a unique sensory terminal region of the spinal cord. Using *in vivo* electrophysiology and calcium imaging, we found that both Krause corpuscle afferent types are A-fiber rapid-adapting low-threshold mechanoreceptors, optimally tuned to dynamic, light touch and mechanical vibrations (40-80 Hz) applied to the clitoris or penis. Optogenetic activation of male Krause corpuscle afferent terminals evoked penile erection, while genetic ablation of Krause corpuscles impaired intromission and ejaculation of males as well as reduced sexual receptivity of females. Thus, Krause corpuscles, which are particularly dense in the clitoris, are vibrotactile sensors crucial for normal sexual behavior.

---

Somatosensory end organs are specialized for the functions of the body region or skin type in which they reside. For example, Meissner corpuscles located in dermal papillae of glabrous skin underlie light touch perception and support fine sensory-motor exchange and dexterity of the hands and digits, while, in hairy skin, longitudinal lanceolate ending complexes associated with hair follicles mediate sensory responses to hair deflection (*1*). Although we have a deep understanding of the somatosensory end organs associated with glabrous and hairy skin, the physiological properties and functions of sensory structures within the mammalian genitalia are not known.

In the late 19th century, Wilhelm Krause first described specialized sensory corpuscles located in human genitalia and other mucocutaneous tissues, including the lips, tongue, and conjunctiva of the eye (*2–5*). He found that corpuscles of the penis and clitoris display either a glomerular shape and contain coiled axons, or they are smaller in size, possess a cylindric shape, and contain simple axonal endings. These sensory structures have been assigned a number of names, including mucocutaneous end-organs (*4*), Krause corpuscles, Krause end bulbs, and genital corpuscles (*6, 7*); we use the name “Krause corpuscles” for these sensory end organs of the male and female genitalia. While the morphological properties of Krause corpuscles were described long ago, their physiological properties and functions have remained a subject of speculation. Here, we describe the anatomical and physiological properties of Krause corpuscle-innervating sensory neurons of the clitoris and penis and their functions in sexual behavior.

## Results

### Morphological diversity and sexually dimorphic density of Krause corpuscles in mouse genitalia

To assess the distribution and density of Krause corpuscles in the genitalia of mice, we stained thick (200 µm) sagittal sections of genital tissue for neurofilament 200 (NF200) to visualize large caliber sensory axons and S100 for terminal Schwann cells, which wrap sensory axon terminals to form corpuscles. In female genitalia, a remarkably high density of Krause corpuscles was observed throughout the clitoris, which is located within the visible protrusion of hairy skin, dorsal to the distal urethra, and between the preputial glands (Fig. 1, A to C, and fig. S1A) (*8*).

**Fig. 1.**
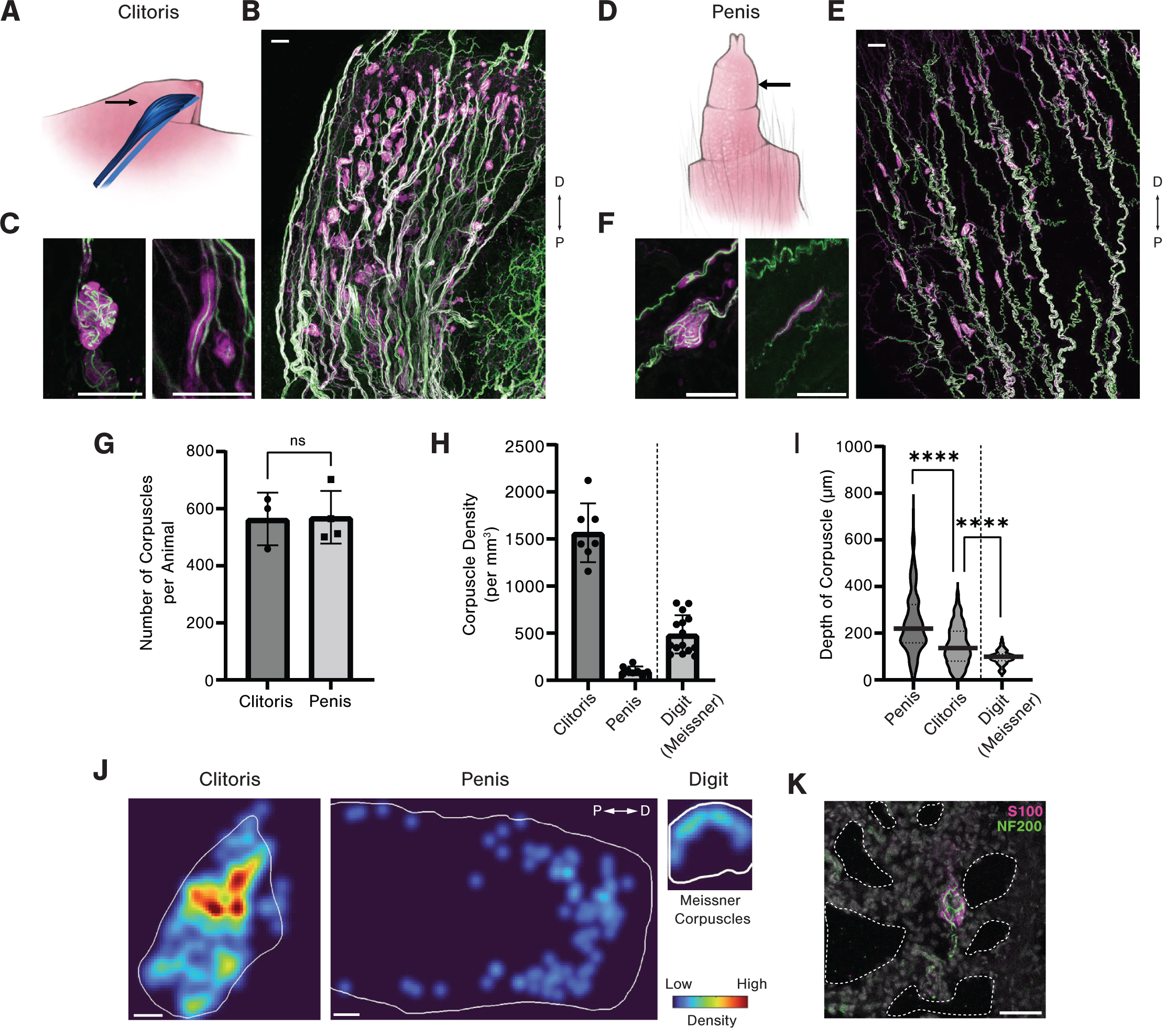
Krause corpuscles are distributed across female and male mouse genitalia. (**A**) Illustration of the mouse clitoris (drawn in blue and indicated by a black arrow) within the visible protrusion of hairy skin. (**B**) Sagittal section (200 µm thick) of the clitoris stained for S100 and NF200. (**C**) Examples of complex (left) and simple (right) Krause corpuscles in the clitoris, stained as in (B). (**D**) Illustration of the glans penis (arrow) externalized from the retracted internal prepuce and hairy external prepuce. (**E**) Sagittal section (200 µm thick) of the glans penis of an adult mouse immunostained as in (B). (**F**) Examples of complex (left) and simple (right) Krause corpuscles in the penis, stained as in (B). (**G**) Total number of corpuscles summed across all sections of clitoris or penis in each animal (n = 3 females [564 mean ± 92.4 standard deviation], 4 males [570 ± 92.3]; p = 0.94; unpaired t-test). (**H**) Density of Krause corpuscles in the mouse clitoris and penis, compared to the density of Meissner corpuscles in the digit. (n = 7 sections from 4 females [1566 corpuscles/mm^3^ ± 311.8], 8 sections from 2 males [100.7 ± 47.1], 15 sections from 2 animals’ digits [486.9 ± 203.1]). (**I**) The depth of corpuscles from the nearest surface of the clitoris, penis, or digit (µm). The violin plot contains 365 corpuscles from 8 sections of penis [243.9 ± 132], 554 from 7 sections of clitoris [146.9 ± 83.3], and 130 from 15 sections of digit [96.3 ± 29.1]. One-way analysis of variance (ANOVA) effect of tissue type (F_2, 1046_= 151.2, p < 0.0001) with post-hoc multiple comparisons test: ****p < 0.0001. In the penis, the corpus cavernosum and adjacent tunica albuginea begin approximately 200 µm below the surface and terminate at 500 µm, as shown in fig. S1B. (**J**) Example heatmap of corpuscle density across a clitoris, penis, and digit. Heatmaps are normalized equally across the tissue types to the densest region of the clitoris. (**K**) Example of a Krause corpuscle adjacent to expanded cavernous space (dotted line) in a penis fixed in the erect state and stained as in (B). Scale bars are all 50 µm, except for the scale bar in J, which is 200 µm. “D” and “P” indicate “distal” and “proximal”, respectively.

Notably, these end organ structures were absent from vaginal tissue (fig. S1D). In male genitalia, corpuscles were observed throughout the glans penis (Fig. 1, D to F) and the internal prepuce, which is a thin sheath covering the glans (fig. S1C) (*9*). While earlier reports estimated clitoral and penile sensory neuron innervation density by measuring the number of nerve fibers entering the genitalia (*10, 11*) or using small fields of view (*12, 13*), we obtained a comprehensive, quantitative assessment of female and male Krause corpuscles by counting the total number of corpuscles across the entire genital tissue (Fig. 1G). Strikingly, despite the different sizes of the female and male genitalia, the total number of Krause corpuscles within the clitoris and penis was comparable, thus resulting in a 15-fold higher density of Krause corpuscles in the clitoris than in the penis (Fig. 1H). For comparison to another highly sensitive skin region, the density of Meissner corpuscles of the digit tips was assessed, revealing three-fold more Krause corpuscles per unit volume of the clitoris than Meissner corpuscles of digit skin (Fig. 1, H and J).

Morphologically, Krause corpuscles of the mouse genitalia spanned a range of shapes and sizes (Fig. 1 C and F, fig. S2, A and B). At one end of this range, corpuscles contained multiple axons tightly coiled like balls of yarn, resembling Krause’s “glomerular corpuscles” or “end-bulbs”, while, at the other end, corpuscles consisted of one or two linear axons, resembling Krause’s “cylindrical,” “simple lamellar,” or Golgi–Mazzoni corpuscles (*7*). We refer to the former as “complex Krause corpuscles” and the latter as “simple Krause corpuscles”. A greater proportion of complex Krause corpuscles was observed in clitoral tissue (93 ± 5% of corpuscles) compared to penile tissue (70 ± 7%) (fig. S1E). Moreover, the clitoris uniquely contained a population of elaborate, lobulated Krause corpuscles with multiple clusters of S100^+^ cells (fig. S2C). Despite a range in the number of axon profiles and terminal complexity across the simple and complex Krause corpuscles, electron microscopy analysis showed that axons in both types of corpuscle were concentrically wrapped by lamellar processes with a variable number of layers (fig. S3), as previously described for Krause corpuscles in the rat (*14*) and human genitalia (*12*).

Unlike Meissner corpuscles, which strictly reside within the dermal papillae, Krause corpuscles were broadly distributed across both the clitoris and penis, often forming along axon bundles with NF200^+^ fibers on opposing sides of the structure (Fig. 1, C, F, J, fig. S2, A and B). In the penis, an enrichment of corpuscles within erectile tissue (corpus cavernosum) was also observed (Fig. 1J). In fact, penile tissue prepared in the erect state displayed corpuscles directly adjacent to expanded cavernous spaces (*14*), lined by CD31^+^ endothelial cells (Fig. 1K and fig. S1F), potentially rendering them responsive to vascular engorgement or pressure changes during erection (*15*). Altogether, these findings establish the presence of morphologically diverse Krause corpuscles within the mouse genitalia, the structural similarity between mouse and human Krause corpuscles, and the remarkably high density of complex Krause corpuscles within the clitoris.

### Krause corpuscles are innervated by TrkB^+^ and Ret^+^ afferents that exhibit sexually dimorphic terminal fields

The physiological properties and functions of Krause corpuscles remain unknown despite their discovery over 160 years ago (*2*). Therefore, we sought mouse genetic tools that enable in-depth morphological analysis, targeted physiological recordings, and functional investigation of Krause corpuscle neurons. An initial survey of mouse genetic tools revealed that two alleles, *TrkB^CreER^*and *Ret^CreER^* (*16, 17*), efficiently labeled NF200^+^ Krause corpuscle neurons with high specificity in both female and male genitalia. *TrkB^CreER^*(tamoxifen at P5) labeled dorsal root ganglion (DRG) sensory neuron axons that terminate in nearly all Krause corpuscles of both the clitoris and penis (Fig. 2A, and fig. S2, A and B, and S4A), and it did not label other, non-Krause corpuscle, axonal endings in genital tissue. These TrkB^+^ axons formed both coiled terminals within complex Krause corpuscles and linear terminals within singly innervated, simple Krause corpuscles (fig. S2, A and B). In contrast, Ret^+^ DRG neuron axons, labeled using the *Ret^CreER^*allele (tamoxifen at E11.5/ E12.5) or the *Ret^CFP^* allele combined with NF200 staining, innervated most Krause corpuscles and were accompanied by additional Ret^-^/NF200^+^ axons (Fig. 2B and fig. S4A). These findings raised the possibility that complex Krause corpuscles are dually innervated by TrkB^+^ and Ret^+^ DRG neurons. To directly test this, we used *TrkB^CreER^; R26^LSL-tdTomato^; Ret^CFP^* mice to simultaneously visualize axonal endings of the TrkB^+^ and Ret^+^ DRG neuron populations, which are largely nonoverlapping (fig. S4B). These double labeling experiments revealed that complex Krause corpuscles contained extensively coiled TrkB^+^ axons and less branched, more peripherally localized Ret^+^ axons, while simple Krause corpuscles contained linear TrkB^+^ axons but lacked Ret^+^ axons (Fig. 2, A and B, and fig. S4C). While this dual-innervation pattern of Krause corpuscles is reminiscent of Meissner corpuscles in glabrous skin (*18*), Krause corpuscles exhibited distinct axonal coiling and distribution patterns (Fig. 1, I to K, fig. S2). Also similar to Meissner corpuscles (*18*), TrkB signaling in DRG sensory neurons is essential for Krause corpuscle formation, as Krause corpuscles were nearly absent in both the clitoris and penis in mice lacking TrkB in sensory neurons (*Avil^Cre^; TrkB^flox/flox^* mice, referred to as *TrkB^cKO^* mice) (Fig. 2C and fig. S5).

**Fig. 2.**
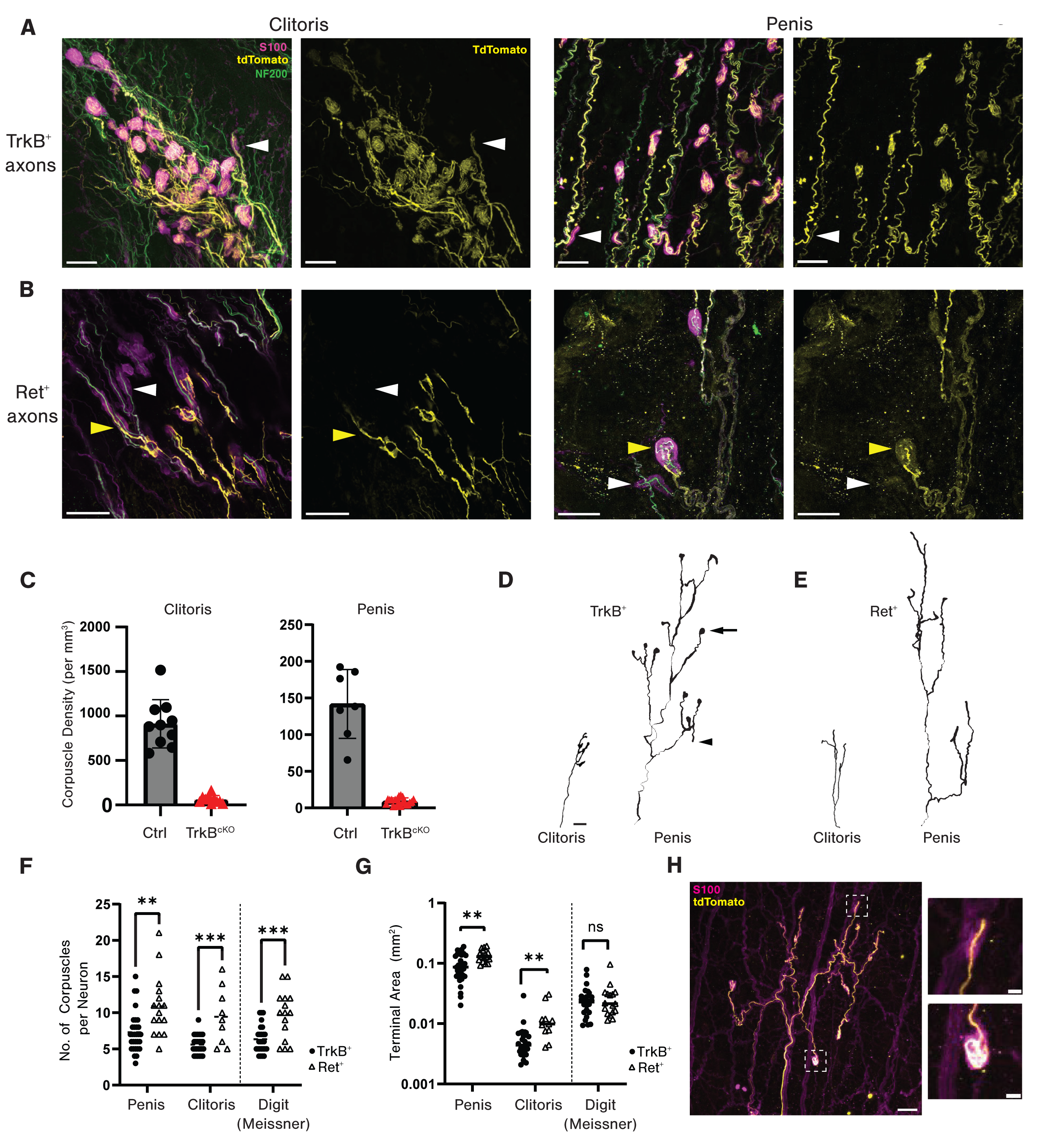
Krause corpuscles are innervated by TrkB^+^ and Ret^+^ afferents with sexually dimorphic terminal fields. (**A**) Representative images of Krause corpuscle afferents in the clitoris (left) and penis (right) labeled in *TrkB^CreER^; Avil^FlpO^; R26^FSF-LSL-Tdtamato^* mice treated with tamoxifen (TAM) at P5. Sections were stained for S100, NF200, and mCherry (tdTomato). TrkB^+^ axons innervate both complex and simple Krause corpuscles (white arrowhead). (**B**) Example of Krause corpuscles afferents labeled in *Ret^CreER^; Avil^FlpO^; R26^FSF-LSL-Tdtamato^* mice treated with TAM at E11.5 or E12.5. Ret^+^ axons are seen alongside additional Ret^-^/NF200^+^ axons in complex corpuscles (yellow arrowhead) and are absent from simple corpuscles (white arrowhead). (**C**) Density of corpuscles in the clitoris and penis of *TrkB^flox/flox^* (Ctrl) and *Avil^Cre^; TrkB^flox/flox^* (TrkB^cKO^) mice (10 sections from three ctrl females [913.4 mean ± 271.1 standard deviation], 10 sections from three TrkB^cKO^ females [62.3 ± 44.5], seven sections from two ctrl males [141.8 ± 46.9], and 20 sections from four TrkB^cKO^ males [8.9 ± 4.7]). (**D** and **E**) Reconstructed traces of single axons in the clitoris and penis labeled in *TrkB^CreER^; Brn3a^cKOAP^* mice (TAM 0.002 mg delivered at P5) (D), and in *Ret^CreER^; Brn3a^cKOAP^* mice (TAM 0.5 mg at E12.5) (E). The AP signal in TrkB^+^ axons often spread into bulbous endings (arrow) in many terminals and more linear endings in others (arrowhead). (**F**) The number of Krause corpuscles innervated by single TrkB^+^ (circle) or Ret^+^ (triangle) afferents in the penis and clitoris, or Meissner corpuscles in digits of the paw, observed with sparse AP labeling (32 TrkB^+^ axons from five males [7.4 ± 2.8], 16 Ret^+^ axons from three males [10.4 ± 4.4], 26 TrkB^+^ axons from four females [5.6 ± 1.3], 11 Ret^+^ axons from three females [9.5 ± 3.6], 26 TrkB^+^ axons from four animals’ digits [6.3 ± 1.9], and 17 Ret^+^ axons from four animals’ digits [9.5 ± 3.4]). Two-way ANOVA revealed no effect of tissue type within TrkB^+^ or Ret^+^ neurons (F_1, 116_= 2.9, p = 0.059) with multiple unpaired t-tests between TrkB^+^ and Ret^+^ neurons of each tissue type: **p < 0.01, ***p < 0.001. (**G**) The area encompassed by the terminals of individual TrkB^+^ or Ret^+^ afferents of the penis [0.095 ± 0.046mm^2^ TrkB and 0.136 ± 0.032mm^2^ Ret], clitoris [0.0058 ± 0.0052 TrkB and 0.012 ± 0.0086mm^2^ Ret], and digit [0.026 ± 0.016mm^2^ TrkB and 0.027 ± 0.020mm^2^ Ret], plotted on a log 10 scale with multiple unpaired t-tests: **p < 0.01. Data were obtained from the same number of sections and animals as in (F). (**H**) Example of a single TrkB^+^ axon, sparsely labeled in a *TrkB^CreER^; Avil^FlpO^; R26^FSF-LSL-Tdtamato^*mouse, terminating in both simple (top inset) and complex (bottom inset) Krause corpuscles. Scale bars from (A) to (H) are 50 µm, and the scale bars of the inset in (H) is 10 µm.

We also visualized axonal arborization patterns of individual TrkB^+^ and Ret^+^ Krause corpuscle afferents using sparse genetic labeling and whole-mount alkaline phosphatase (AP) staining of genital tissue (Fig. 2, D and E). In both the clitoris and the penis, individual Ret^+^ DRG neurons innervated a greater number of corpuscles and covered a larger terminal area than TrkB^+^ neurons (Fig. 2, F and G). Furthermore, the terminal innervation areas of individual TrkB^+^ and Ret^+^ DRG neurons were 11 and 16 times smaller, respectively, in the clitoris compared to the penis (Fig. 2G), despite these neurons forming a similar number of corpuscles (Fig. 2F). This finding is aligned with the 15-fold higher density of Krause corpuscles observed in the clitoris than in the penis (Fig. 1H). Additionally, we observed that the terminals from an individual TrkB^+^ neuron could include both bulbous and linear endings (Fig. 2, D and H), indicating that a single TrkB^+^ neuron can form both types of Krause corpuscle. This diversity of terminal structures associated with individual Krause corpuscle afferents may endow them with a range of sensitivities or tuning properties.

In addition to Krause corpuscle-associated neurons, we observed free nerve endings formed by other DRG sensory neuron subtypes in the genitalia, including CGRP^+^ fibers, MRGPRD^+^ fibers, and NF200^+^ fibers that are not corpuscle-associated. These free nerve endings were observed throughout the genital tissue, often terminated close to the surface of the tissue, and emerged from axons that occasionally passed through Krause corpuscles (fig. S6). It is noteworthy that Merkel cells, which associate with slowly adapting low-threshold mechanoreceptors (*19*), were absent from genital tissue, although they were observed in abundance in adjacent hairy skin (fig. S6E). Therefore, while a range of DRG neuron subtypes innervate the genitalia, TrkB^+^ and Ret^+^ DRG sensory neurons uniquely form Krause corpuscles.

### Krause corpuscle afferents project from the genitalia to the dorsomedial spinal cord

We next examined the central termination patterns of the TrkB^+^ and Ret^+^ DRG neurons that innervate Krause corpuscles. In initial experiments, CTB tracers conjugated to different fluorophores were injected into genital tissues and, for comparison, the adjacent mid-line hairy skin (Fig. 3, A and B) to label anatomically distinct populations of sensory neurons whose cell bodies reside within bilateral L6 and S1 DRGs, but not within the nodose ganglia (fig. S7, A and B). CTB-labeled cell bodies of DRG neurons that innervate the genitalia and adjacent hairy skin regions were almost entirely non-overlapping (fig. S7A). The central terminals of sensory neurons innervating the genital tissue and adjacent hairy skin were observed in the lower lumbar and upper sacral spinal cord in highly segregated patterns (Fig. 3, A and B). While sensory neurons innervating midline hairy skin terminated within medial regions of both the left and right spinal cord dorsal horn, genital-innervating sensory neurons terminated in a unique, midline region of the spinal cord, located between the dorsal column and central canal; this region is often called the dorsal gray commissure (DGC) (*20–22*).

**Fig. 3.**
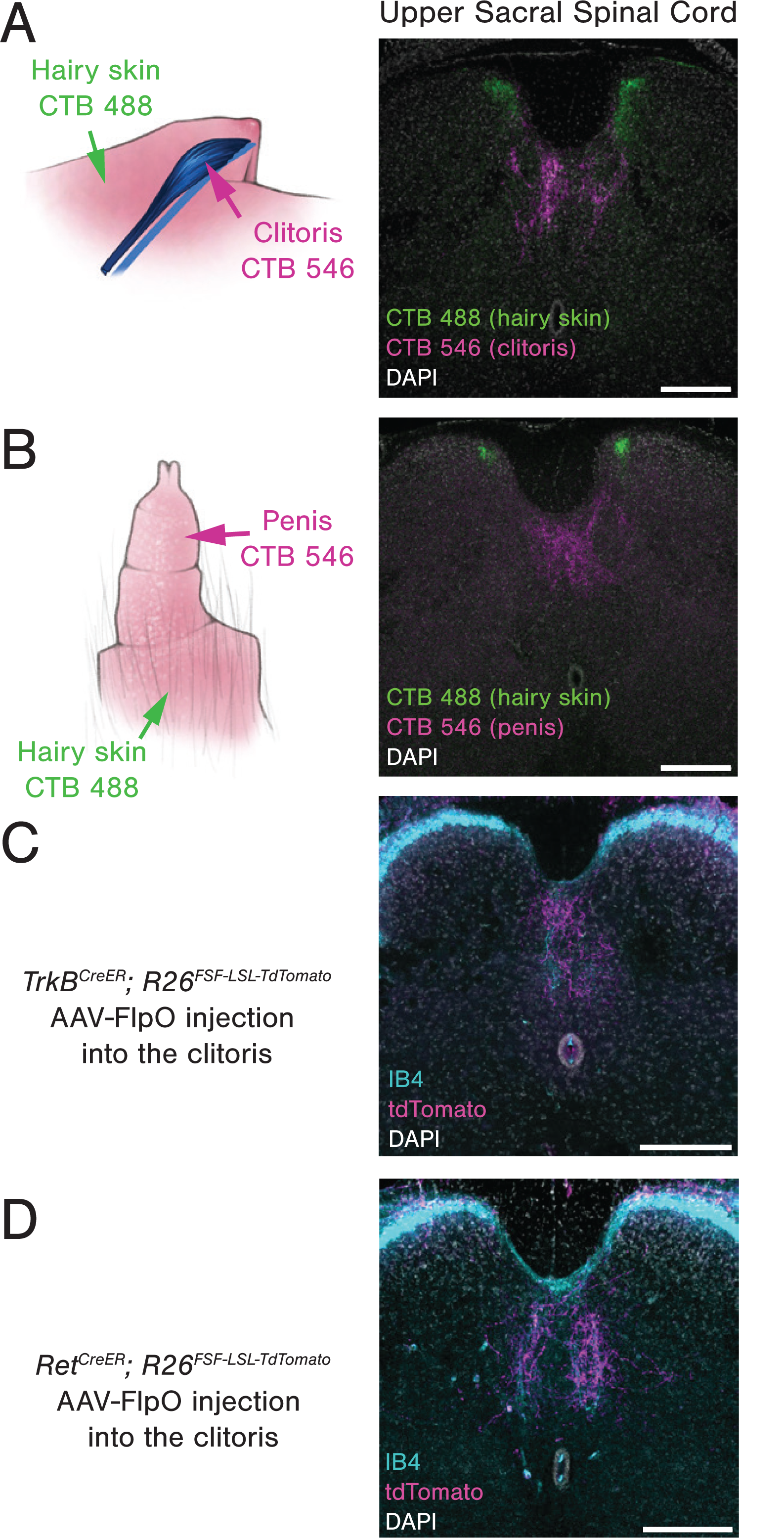
Krause corpuscle afferents project to the dorsomedial spinal cord. (**A** and **B**) Retrograde labeling of DRG neurons innervating the clitoris (A) or penis (B) and the adjacent midline hairy skin region using CTB tracers conjugated to different fluorophores (hairy: CTB-488; genital: CTB-546). Coronal sections of the upper sacral spinal cord are shown on the right. (**C** and **D**) Labeling of Krause corpuscle afferents using injection of AAV2-retro-hSyn-FlpO into the clitoris of *TrkB^CreER^; R26^FSF-LSL-Tdtamato^* (TAM 0.5 mg at P5) (C) or *Ret^CreER^; R26^FSF-LSL-Tdtamato^* (TAM 3 mg at E12.5) (D) animals, with IB4 counterstaining to label the nonpeptidergic nociceptors. Scale bar in A-D is 200 µm.

To specifically visualize the spinal cord termination patterns of TrkB^+^ and Ret^+^ Krause corpuscle afferents, we injected AAV-FlpO into the genital tissue of *TrkB^CreER^*or *Ret^CreER^* mice harboring either a dual-recombinase-dependent fluorescent reporter allele (*R26^FSF-LSL-tdTomato^*) or a dual-recombinase-dependent AP reporter allele (*Tau^FSF-iAP^*). As observed in the CTB pan-neuronal labeling experiments, TrkB^+^ and Ret^+^ afferents innervating the clitoris terminated exclusively in the DGC region of the spinal cord (Fig. 3, C and D, fig. S7C). Moreover, whole mount AP labeling experiments revealed that axons of individual TrkB^+^ Krause corpuscle afferents bifurcated upon entering the spinal cord and formed clusters of collaterals extending along the rostro-caudal axis (fig. S7C). While collaterals were enriched between L6 to S1, some collaterals containing fewer terminal branches extended a few segments away but did not reach upper lumbar spinal segments or the brainstem. Thus, Krause corpuscle afferents form an elaborate pattern of axon terminals within the DGC region of the lumbosacral spinal cord.

### Krause corpuscle afferents are fast conducting low-threshold mechanosensory neurons optimally tuned to mechanical vibration

Genetic access to Krause corpuscle afferents allowed us to address long-standing questions of their physiological properties and functions. Therefore, we developed a preparation for direct mechanical and thermal stimulation of the external genitalia during *in vivo* electrophysiological measurements and calcium imaging of cell bodies of TrkB^+^ and Ret^+^ Krause corpuscle-innervating neurons within L6 DRGs (Fig. 4, A and F). Using *in vivo* multi-electrode array (MEA) recordings of L6 DRG neurons, TrkB^+^ Krause corpuscle afferents were optotagged using *TrkB^CreER^*; *Avil^Flpo^*; *R26^FSF-LSL-ReaChR^* mice (Fig. 4, A and B). In male mice, TrkB^+^ Krause corpuscle neurons were activated using a combination of mechanical and optogenetic stimuli applied to the glans penis to establish their receptive field locations. Male TrkB^+^ Krause corpuscle afferents exhibited conduction velocities characteristic of A fibers (3-11 m/s based on their latency to optogenetic stimuli, n = 4) (*1*) and robust activation by light touch of the penis (Fig. 4, B and C). These neurons exhibited low mechanical thresholds (1-10 mN), rapid adaptation (RA), firing upon the onset and offset of step indentations (Fig. 4D), and precise phase locking to each cycle of mechanical vibrations up to 120Hz, the highest frequency tested (Fig. 4E).

**Fig. 4.**
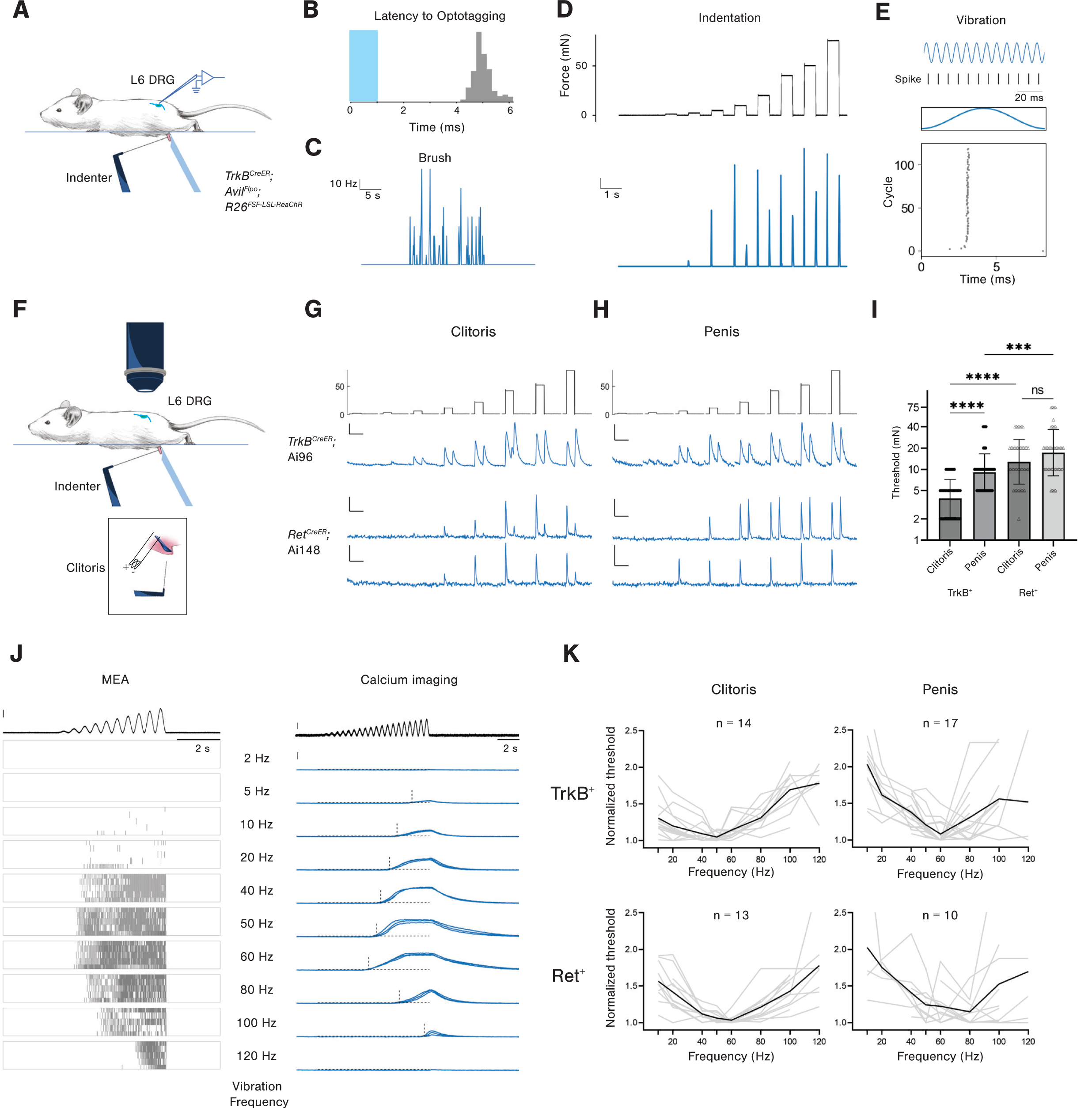
TrkB^+^ and Ret^+^ Krause corpuscle afferents are fast-conducting, low-threshold mechanical vibration sensors. (**A**) Schematic of mechanical stimulation of the glans penis during *in vivo* MEA recording of L6 DRG neurons. The blue object on the opposite side of the indenter stabilizes the glans penis. (**B**) Latency of spikes recorded in the DRG after optogenetic activation (1 ms pulse, shown in blue) of a TrkB^+^ Krause corpuscle neuron of the penis. (**C-E**) The TrkB^+^ neuron optotagged in (B) exhibits a robust response to brushing the penis (C), a rapid-adapting response to step indentations ranging from 1 mN to 75 mN (D), and phase-locking to sinewave vibration stimuli (120 Hz, 19 mN) across 120 cycles (E). (**F**) Schematic of *in vivo* calcium imaging of L6 DRG neurons. In male mice, mechanical stimulation was applied as shown in (A). In female mice, electrical stimulation was applied to the exposed dorsal nerve of clitoris after mechanical stimulation of the clitoris (inset). (**G** and **H**) Representative calcium signals from TrkB^+^ neurons (labeled in *TrkB^CreER^*; Ai96 mice, TAM 0.5 mg delivered at P5) and Ret^+^ neurons (labeled in *Ret^CreER^*; Ai148 mice, TAM 3mg delivered at E12) following step indentations of the clitoris (G) and penis (H). Scale bar in (G and H): 5% ΔF/F and 5 seconds. All TrkB+ neurons (n=13 innervating the clitoris, n=16 innervating the penis) and a subset of Ret^+^ neurons (9 of 18 innervating the clitoris, 8 of 10 innervating the penis) showed an ON-OFF response, while the remaining Ret^+^ neurons showed only an ON response (bottom trace). (**I**) Comparison of indentation thresholds (log 2 scale) of TrkB^+^ and Ret^+^ Krause afferents of the clitoris and penis, based on calcium imaging results in (G and H) (two-way ANOVA, ***p < 0.001, and ****p < 0.0001). (**J**) Representative responses of TrkB^+^ Krause corpuscle afferents to ramping-force vibration stimuli of different frequencies. Left: the top black trace represents the ramping vibration stimuli applied during MEA recordings (0 to 20 mN at frequencies varying from 2 Hz to 120 Hz, vertical scale bar: 5 mN); the raster plots below show the spikes from a representative TrkB^+^ Krause afferent innervating the penis across five trials. Right: the top black trace represents the ramping-force vibration stimuli applied during calcium imaging (vertical scale bar: 2 mN); the blue traces below show the calcium signals (vertical scale bar: 100% ΔF/F) from an example TrkB^+^ Krause afferent innervating the penis across three trials. The horizontal dashed lines indicate the threshold for response (five times the standard deviation of baseline), while the vertical dashed lines indicate the timing of response onset. (**K**) Normalized frequency turning curves. The mechanical threshold for each vibration frequency is normalized to the minimum threshold of the neuron. Curves from individual neurons are shown in gray, and the average curve is shown in black. N indicates the number of neurons.

Because of the low throughput of the electrophysiological recordings, we complemented the electrophysiological analysis with *in vivo* calcium imaging experiments using both male and female mice that express GCaMP6 in the TrkB^+^ or Ret^+^ DRG sensory neuron populations (Fig. 4F). Precise mechanical stimuli applied to the glans penis allowed identification of TrkB^+^ or Ret^+^ Krause corpuscle afferents of males, while electrical stimulation of the dorsal nerve of the clitoris (*23*) was used to distinguish clitoris-innervating afferents from neurons that innervate the overlying hairy skin of females (fig. S8A). Consistent with findings from the *in vivo* MEA electrophysiological recordings in the male, these calcium imaging experiments revealed that both penis- and clitoris-innervating TrkB^+^ neurons are RA low-threshold mechanoreceptors (LTMRs), exhibiting both an ON and an OFF response to step indentations of the genitals (Fig. 4, G and H). This analysis also showed that TrkB^+^ sensory neurons innervating the clitoris are more sensitive than those innervating the penis (Fig. 4I). In addition, as a population, the Ret^+^ Krause corpuscle afferents in both the penis and clitoris exhibited higher mechanical force thresholds than the TrkB^+^ afferents, and a subset of Ret^+^ neurons exhibited responses only to the onset of indentation (Fig. 4, G to I). The distinct mechanical thresholds and adaptation properties of TrkB^+^ and Ret^+^ Krause afferents may reflect the observed differences in their terminal morphology within corpuscles (Fig. 2).

As RA-LTMRs that innervate other skin regions such as glabrous skin of the digits display preferred sensitivity to certain vibration frequencies (*1*), we next assessed force thresholds of the two Krause corpuscle afferent types across a range of vibration frequencies. For both the clitoris and penis, TrkB^+^ and Ret^+^ Krause corpuscle neurons exhibited the lowest force thresholds in response to 40-80 Hz vibration stimuli, indicating exquisite sensitivity to vibrotactile stimuli (Fig. 4, J and K). The mechanotransduction channel Piezo2 was highly localized to axons within, but not outside, the Krause corpuscle, and not to lamellar cells, in both the penis and clitoris (fig. S8B) (*24*), providing a molecular basis for the mechanosensitivity of Krause corpuscle afferents (*25, 26*). Additionally, early literature had proposed that Krause corpuscles are thermoreceptors based on anatomical considerations (*27, 28*), a notion that has persisted in contemporary texts and reviews (*6, 29*). However, we observed that TrkB^+^ Krause corpuscle afferents are insensitive to changes in temperature (fig. S8C). Thus, TrkB^+^ and Ret^+^ Krause corpuscle afferents are A-fiber RA-LTMRs, exhibiting distinct mechanical thresholds and optimal sensitivity to 40-80 Hz mechanical vibrations, and TrkB^+^ Krause corpuscle afferents of the clitoris are the most sensitive of the genital-innervating RA-LTMRs.

### Functions of Krause Corpuscles in sexual behaviors

The high density and exquisite vibrotactile sensitivity of Krause corpuscle afferents, and the availability of genetic tools to study them, prompted us to ask whether Krause corpuscles contribute to sexual behavior. We began by testing whether activation of Krause corpuscle afferents is sufficient to evoke sexual reflex behaviors. Previous studies have reported that restrained male mice can only display mechanically evoked sexual reflexes after spinal cord transection to eliminate descending inhibitory signals (*30, 31*). Following spinal transection at T9, male mice displayed robust erectile responses to brushing and 50 Hz mechanical vibration applied to the penis in awake, non-anesthetized animals (Fig. 5B and movie S1). These responses typically began with an erection, characterized by distension and reddening of the penis, often followed by “cup”, a flared full engorgement with blood, and, in many cases, “flip”, a brief dorsiflexion of the penis (Fig. 5A) (*30*). Because Krause corpuscle afferents were maximally activated by brush and mechanical vibrations (Fig. 4), we tested whether optogenetic activation of Krause corpuscle afferent terminals in the penis can recapitulate mechanically evoked reflex responses. We found that direct optogenetic stimulation of the penis (10 Hz, 2 ms pulse for 20 s) of *TrkB^CreER^; Avil^Flpo^; R26^FSF-LSL-ReaChR^* mice, which express ReaChR in TrkB^+^ Krause corpuscle afferents, led to erectile responses in 6 out of 10 animals. In contrast, light pulses did not evoke erection when delivered to the penis of control animals lacking opsin (Fig. 5C). Moreover, optical stimulation of the glans penis of mice expressing a faster opsin, CatCh (*32*), in TrkB^+^ Krause corpuscle afferents using a higher frequency stimulus (20Hz, 1ms pulse for 20s) led to erectile responses in 5 out of 5 animals tested (Fig. 5C and movie S2). Reflex responses to optogenetic stimulation of Ret^+^ Krause corpuscle afferents were not tested, because, in addition to Ret^+^ DRG sensory neurons, the *Ret^CreER^* allele also labels autonomic fibers (*33*), a confounding issue when assessing sexual reflexes. Nevertheless, these findings indicate that optogenetic activation of TrkB^+^ Krause corpuscle afferents is sufficient to initiate sexual reflexes in male mice.

**Fig. 5.**
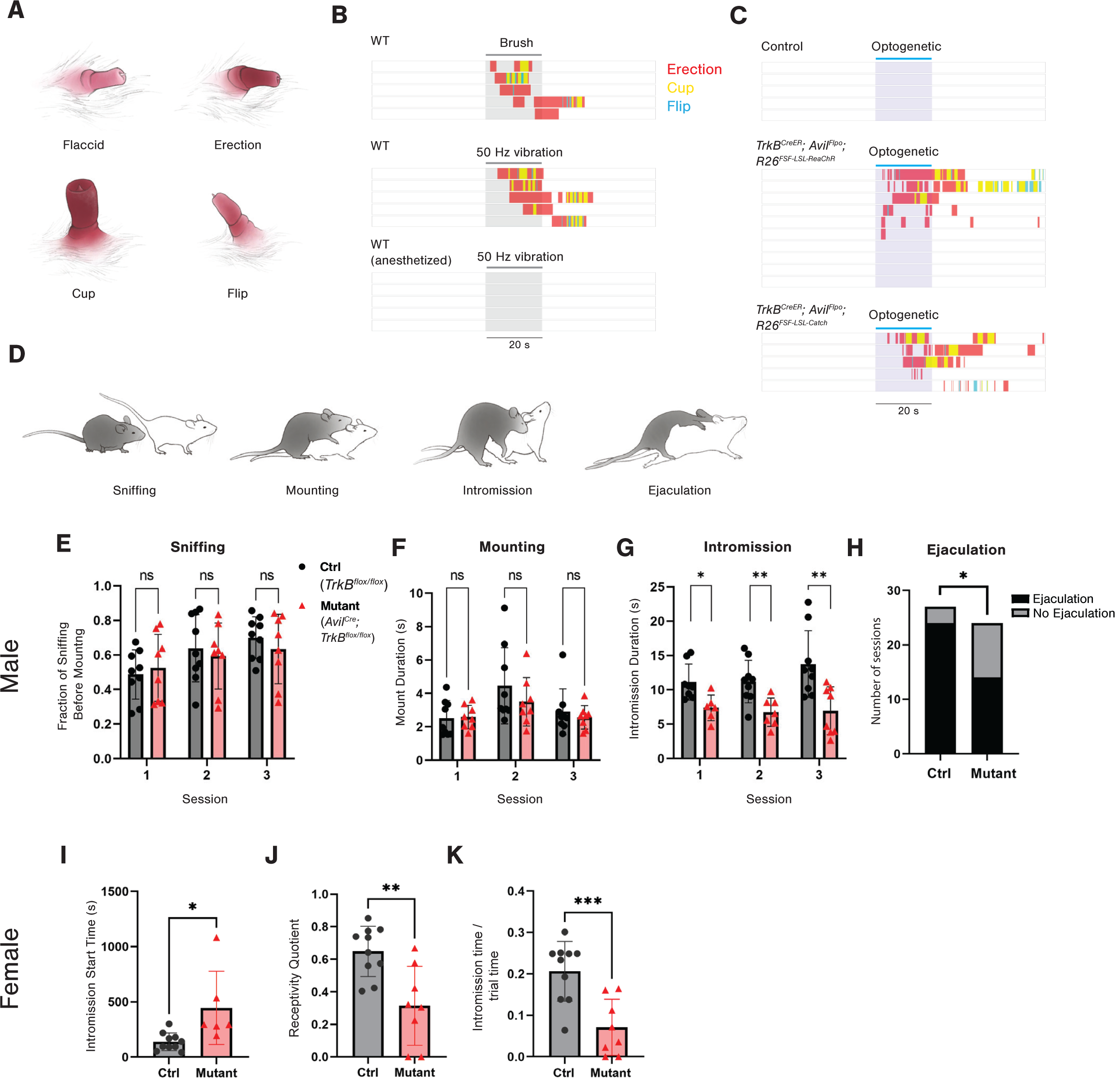
Krause corpuscles mediate normal sexual behaviors. (**A**) Illustrations of the four states of the mouse penis during sexual reflexes. (**B**) Ethograms depicting the reflex responses of awake, spinalized mice to brush stimulation of the glans penis (top), 50 Hz vibration of the glans penis (middle), and 50 Hz vibration of the glans penis during general anesthesia using isofluorane (bottom). (**C**) Ethograms showing sexual reflex responses to optogenetic stimulation of the penis of mice without opsin expression (top), mice with expression of ReaChR in TrkB^+^ sensory neurons (middle), and mice with expression of CatCh (Ai80) in TrkB^+^ sensory neurons (bottom). (**D**) Schematics of mouse mating behaviors. The male mouse is colored dark gray, and the female is colored white (For experiments, only animals with black coats were used). (**E** to **G**) Comparisons of the mating behavior of primed, receptive wild-type females with control (n = 9) and *TrkB^cKO^*(n = 8) males: quantification of sniffing (E), average duration per mounting bout (F), and average duration per intromission bout (G) across three sessions that were at least one week apart. Multiple unpaired t-test, *p < 0.05, **p < 0.01. (**H**) Comparison of the number of sessions with successful ejaculation summed over the three sessions between controls and *TrkB^cKO^*animals. Of the 27 trials for controls, 24 resulted in successful ejaculations within 75 minutes, while of the 24 trials for mutants, only 14 achieved successful ejaculations within 75 minutes. Fisher’s exact test, p = 0.022. (**I** to **K**) Comparisons of the mating behavior of naturally cycling, experienced control (n = 10) and *TrkB^cKO^* (n = 8) females: start time of intromission (I), receptivity quotient (calculated as the total intromission time divided by the sum of the total intromission time and mounting time) (J), and intromission time divided by total trial time (K). Unpaired t-test, *p < 0.05, **p < 0.01, ***p < 0.001.

To determine the necessity of Krause corpuscles in sexual behaviors, we used *TrkB^cKO^* mice, which lack Krause corpuscles (Fig. 2C). We first assessed vibration-induced reflexes of control and *TrkB^cKO^* animals, finding no clear deficiency in mechanically evoked erectile responses in the mutants (fig. S9A). This finding suggests that either Krause corpuscles are not necessary for mechanically evoked sexual reflexes in surgically spinalized mice or that Krause corpuscle afferents remain present in the mutants even though they are mostly devoid of mature corpuscles. Therefore, we next explored the possibility that Krause corpuscles contribute to mating behaviors of awake, normally behaving mice using male and female *TrkB^cKO^* animals without the confounds of spinalization (Fig. 5D and movie S3).

We found that *TrkB^cKO^* males lacking Krause corpuscles were motivated to mate to a comparable extent as control littermates, indicated by a similar percentage of time that *TrkB^cKO^* and control male mice spent exploring and sniffing wildtype, hormonally primed females (Fig. 5E). *TrkB^cKO^* males also displayed a normal mounting motion (Fig. 5F), suggesting that general motor coordination is largely normal in the mutants, consistent with previous gait analysis of *TrkB^cKO^* mice (*18*). However, *TrkB^cKO^* males exhibited impaired intromission, the stage of mating when the penis is inserted into the vagina, observable as rhythmic thrusts. Compared to controls, *TrkB^cKO^* males displayed shorter bouts of intromission, a delayed start of intromission, a reduced total amount of intromission time, and a trend toward increased inter-intromission intervals over successive trials (Fig. 5G and fig. S9, B to H), suggestive of aberrant sensory feedback from the penis. Importantly, fewer *TrkB^cKO^*males achieved ejaculation within the 75-minute session, compared to controls (Fig. 5H). Thus, Krause corpuscles of the penis are necessary for normal intromission and ejaculation.

We also assessed mating behaviors of *TrkB^cKO^* females, which lack Krause corpuscles of the clitoris. Interestingly, while no differences were observed in the performance of gonadally intact, inexperienced mutant females that were hormonally primed across any of these behavioral measures (fig. S10, A to F), mating behaviors of experienced females while naturally cycling during proestrus or estrus were markedly aberrant. We observed a longer latency for receptivity to intromission, a reduced proportion of males’ mounting attempts that result in successful intromission (intromission quotient), and less total intromission time between *TrkB^cKO^* females and wildtype males, compared to control females and wildtype males (Fig. 5, I to K, and fig. S10, G and H). Thus, loss of the dense population of Krause corpuscles in the clitoris leads to reduced sexual receptivity of experienced female mice.

## Conclusions

Our findings show that Krause corpuscle afferents of the mouse genitalia are low-threshold, rapidly adapting mechanoreceptors optimally sensitive to 40-80 Hz mechanical vibrations, which is remarkably comparable to frequencies used in vibrating devices for human sexual stimulation (*34*). We speculate that low force amplitude microvibrations associated with dynamic skin-skin contact during the motion of intromission activates Krause corpuscle afferents in both male and female genitalia. It is also noteworthy that the vibrotactile signals emanating from Krause corpuscles are conveyed to a unique region of the dorsal-medial spinal cord, just caudal to the spinal ejaculation generator which modulates parasympathetic and sympathetic preganglionic neurons and pudendal motoneurons that control erection and ejaculation (*21, 35–38*). Thus, while other DRG neuron populations may also contribute to sexual behaviors (*39*), Krause corpuscle afferents are optimally tuned to vibratory stimuli and transmit vibrotactile information from the genitalia to the DGC region of the spinal cord to engage the spinal cord sexual reflex circuitry.

Whole mount imaging of Krause corpuscles across the clitoris and penis revealed a comparable number of these vibrotactile end organs in the male and female genitalia, however the clitoris has a remarkably high corpuscle density due to its much smaller size. This intriguing observation suggests the existence of a common innervation pattern of the penis and clitoris during early stages of genital development, followed by divergent genital tissue growth during later development, thereby leading to a highly sexually dimorphic density of Krause corpuscles in adulthood.

Finally, our functional experiments show that vibrotactile information encoded by Krause corpuscle afferents can evoke penile erection to support proper intromission and successful ejaculation. In female mice, Krause corpuscle afferents are crucial for normal mating drive or receptivity. These findings suggest that, in addition to their roles in initiating genital reflexes, Krause corpuscle afferents may also mediate the positive valence associated with vibrotactile stimulation of the genitalia.

## Acknowledgments

We thank Ginty lab members for discussions and comments on the manuscript, O. Mazor and P. Gorelik (HMS Research Instrumentation Core) and J. LeBlanc (HMS Neurobiology Machine Shop) for help with design and construction of the physiology setup and stimulation devices, and A. Emanuel for code used for the MEA data analysis. We thank K. McKenna for discussions about sexual reflex measurements, S. Lima and S. X. Zhang for advice on mating behaviors, and W. Luo for discussions during early stages of this work.

## Funding

Stuart H.Q. and Victoria Quan fellowship (LQ)

Northeastern University Office of Undergraduate Research and Fellowships (MI) Howard Hughes Medical Institute–Jane Coffin Childs Fellowship (A.H.)

NIH grant NS097344 (DDG) NIH grant AT011447 (DDG)

The Bertarelli Foundation (DDG)

The Hock E. Tan and Lisa Yang Center for Autism Research (DDG)

The Lefler Center for Neurodegenerative Disorders (DDG) Howard Hughes Medical Institute (DDG)

## Author contributions

Conceptualization: LQ, MI, DDG Methodology: LQ, MI, AH Investigation: LQ, MI Visualization: LQ, MI

Funding acquisition: LQ, MI, AH, DDG Supervision: LQ, DDG

Writing – original draft: LQ, MI, DDG

Writing – review & editing: LQ, MI, AH, DDG

## Data and materials availability

All data presented in the manuscript or the supplementary materials are available upon request from the corresponding author. The mouse strains used in this study are available from the corresponding author under material transfer agreement with Harvard University.

## Supplementary Materials

Materials and Methods

Figs. S1 to S10

Movies S1 to S3

**Movie S1. Sexual reflexes in response to 50Hz-vibration stimuli.**

**Movie S2. Sexual reflexes in response to optogenetic stimulation of TrkB^+^ Krause afferents, labeled by *TrkB^CreER^; Avil^Flpo^; R26^FSF-LSL-ReaChR^*.**

**Movie S3. Representative mating behaviors: sniffing, mounting, intromission, and ejaculation.**

## Materials and Methods

### Animals

Animals were handled according to protocols approved by the Harvard Standing Committee on Animal Care and are in accordance with federal guidelines.

### Mouse Lines

All mice used in this study have been previously described, including: *TrkB^CreER^*(JAX 027214) (*16*), *Ret^CreER^* (MGI 4437245) (*17*), *Advillin^Cre^* (JAX 032536) (*40*), *Advillin^FlpO^* (*41*), *Ret^CFP^* (MGI 3777555) (*42*), *TrkB^flox^* (*43*), *Brn3a^cKOAP^* (JAX 010558) (*44*), *Tau^FSF-iAP^* (*45*), *MrgprD^GFP^* (*46*), *PLP^EGFP^* (JAX 033357) (*47*), *Piezo2^smFP-FLAG^* (*48*), *R26^FSF-LSL-ReaChR-mCitrine^* (JAX 024846), Ai14 (*R26^LSL^-^tdTomato^*; JAX 007914) (*49*), Ai65 (*R26^FSF-LSL-tdTomato^*; JAX 021875)(*50*), Ai96 (*R26^LSL-GCaMP6s^*)(*50*), and Ai148 (*TIGRE^LSL-GCaMP6f-tTA2^*) (*51*). All lines were kept on a mixed background, while *Advillin^Cre^* and *TrkB^flox^* were bred from mixed background to C57Bl/6 background for two generations for mating behavior testing. For *Advillin^Cre^*;*TrkB^flox/flox^*animals, non-neuronal recombination in ear or tail was routinely detected, likely due to the leaky expression of *Advillin^Cre^*. However, the observed non-neuronal recombination of the *TrkB^flox^* allele was not a result of germline deletion of TrkB, which is lethal (*52*).

### Tamoxifen Treatments

Tamoxifen was dissolved in 100% ethanol, diluted 1:2 in sunflower seed oil, and vacuum-centrifuged for 30 minutes. The TrkB^+^ population was densely labeled for immunohistochemistry and physiology with an intraperitoneal injection of 0.5mg of tamoxifen at postnatal day 5 (P5) and sparsely labeled for AP and immunohistochemistry with 0.002 - 0.005mg of tamoxifen. In some cases, tamoxifen (3mg) was administrated to the pregnant mother at embryonic day 14.5 or 15.5 (E14.5/E15.5) through oral gavage, to label the same populations (*18*). The Ret^+^ population was densely labeled with 3mg of tamoxifen delivered to the pregnant mother through oral gavage at E11.5 or E12.5 and sparsely with 0.5mg at E12.5.

### Perfusion and Post-fixation

Mice were anesthetized with isoflurane and trans-cardially perfused with approximately 15mL of 1x PBS with heparin (10U/mL) and fixed with approximately 15mL of 4% paraformaldehyde (PFA) in 1x PBS. Once perfused, the vertebral column and brain, if needed, were removed and post-fixed in 4% PFA in PBS at 4°C overnight. In male mice, genital tissue including the external prepuce, was removed at the base of the penis following an incision in the scrotal area. In female mice, the entire perineal area was removed from above the protrusion of hairy skin (homologous to the “external prepuce” in males) to below the vaginal opening. The prostatic urethra, from the bladder to the base of the penis in males, and the vaginal canal in females were roughly dissected. Genital tissue was post-fixed in Zamboni solution (phosphate-buffered picric acid-formaldehyde) at 4°C overnight, and tissues were washed in 1x PBS the following day.

### Fixation of the erection state

The penis was collected after the mouse was freshly perfused with 1x PBS with heparin and the penis was isolated from the prepuces. One suture was fastened around the proximal base of the penis (proximal corpus cavernosum, not the corpus cavernosum glans) and the other was loosely wrapped around the base of the glans, just distal to the 90° bend. A 30 G needle attached to a 3mL syringe containing Zamboni solution or 2% PFA was inserted between the sutures. Care was taken to ensure that the needle stayed inside the glans penis. Gentle pressure was applied to the syringe until the penis assumed a “cup” shape, and, while maintaining pressure on the syringe, the distal suture was fastened. As the syringe was removed, the sutures were further tightened, and the tissue was post-fixed in Zamboni solution at 4°C overnight. Subsequently, the tissue was transferred to 30% sucrose at 4 °C until it sank to the bottom, followed by embedding in OCT for cryosectioning.

### Immunohistochemistry

Spinal cords and DRGs were isolated with forceps following removal of the dorsal and ventral vertebral column, and nodose ganglia were dissected upon removal of the overlying submandibular glands and musculature. Penile tissue was isolated from the preputial glands and external and internal prepuces, and the proximal portion of the penis was discarded. Working proximal to distal, the clitoris was isolated from perineal tissue by removal of remaining pelvic musculature and preputial glands, then by fine dissection of the clitoris and urethra from above the vaginal canal and inside of the external prepuce. Glabrous digit-tips were cut from the paws for staining.

Tissue was cryoprotected in 30% sucrose in 1x PBS at 4°C overnight, embedded in Neg-50, and frozen at −80°C. Spinal cord and DRG samples were sectioned transversely using a cryostat (Leica) at 30 µm and placed onto slides. Genital tissue and glabrous digit-tips were sectioned sagittally at 30 µm thickness and placed onto slides or at 200 µm and placed into a well plate with 1x PBS. Thin cryosections were allowed to dry overnight at room temperature. Sections were rehydrated with 1x PBS, blocked with 5% normal donkey serum in 0.1% PBST (0.1% Triton X-100 in 1x PBS) for two hours at room temperature, and incubated with primary antibody in blocking solution for two days at 4°C. The slides were rinsed in PBST 3×10 minutes and incubated with secondary antibody in blocking solution overnight at 4°C. Slides were then rinsed in PBST 3×10 minutes and mounted with Fluoromount-G. Sections were imaged on a Zeiss LSM 700 or 900 confocal microscope.

Immunohistochemistry of thick sections was performed using previously described protocols for whole-mount tissue (*45*). Sections were washed in 0.3% PBST 5×1 hour and incubated in primary antibody in blocking solution (5% normal donkey serum, 75% PBST, 20% DMSO) at room temperature for 3-5 days under gentle agitation. Following 0.3% PBST washes 5×1 hour, secondary antibody in blocking solution was added for 2-4 days at room temperature. Sections were then washed in 0.3% PBST 2×1 hour, stained with DAPI (1µg/mL, 5 minutes) to visualize nuclei, further washed in 0.3% PBST 3×1 hour, and dehydrated in serial methanol concentrations (50%, 75%, 100%, 1 hour each) and then overnight in 100% methanol. Tissue was cleared in BABB (1:2, Benzyl Alcohol : Benzyl Benzoate) and imaged on a Zeiss LSM 700 confocal, 900 confocal, or AxioZoom stereoscope while fully submerged. Dehydrated tissue was stored in methanol at 4°C.

The primary antibodies used in this work were chicken anti-GFP (Aves Labs, GFP-1020, 1:500), goat anti-GFP (US Biological, G8965-01E, 1:500), goat anti-mCherry (CedarLane, AB0040-200, 1:500), rabbit anti-CGRP (Immunostar, 24112, 1:500), chicken anti-NF200 (Aves Labs, NFH, 1:200), rabbit anti-NF200 (Sigma, N4142-.2ML, 1:500), rabbit anti-S100 (ProteinTech, 15146-1-AP, 1:200-1:500), rat anti-TROMA-1 (DSHB, AB_531826, 1:200), goat anti-CD31 (R&D Systems, AF3628, 1:500), guinea pig anti-FLAG (1:500) (*48*) and IB4 (Alexa 647 conjugated) (Invitrogen, I32450, 1:500).

### Quantification of corpuscle structure and distribution

To determine the total number of corpuscles per animal, care was taken in collecting all of the 200 µm sections from a given clitoris or penis sample and staining them with antibodies for S100 and NF200. All sections were imaged under a confocal microscope. The number of corpuscles in each section was manually counted using ImageJ, with the location of each corpuscle preserved in an ROI file. The borders of imaged tissue were manually outlined and saved. Each section’s volume was computed by multiplying the area of the outlined region by the height of the Z-stack, allowing for calculation of corpuscle density. To generate distribution heatmaps, the ROI files containing the location of each corpuscle and the tissue outline were imported to MATLAB. The distribution of corpuscles was binned in a grid, and a Gaussian filter was used to smooth the density distribution.

“Complex” Krause corpuscles (also called Krause glomerular corpuscles and Krause end bulbs) were defined as Krause corpuscles containing tightly coiled axons. Complex Krause corpuscles often exhibited globular shapes, while some also exhibited elongated shapes with convoluted axonal profiles. “Simple” Krause corpuscles (also called Krause cylindrical corpuscles or simple lamellar corpuscles) were defined as the corpuscles containing 1 or 2 linear (non-coiled) axons. All simple corpuscles have an elongated shape, thereby exhibiting some similarity with Pacinian corpuscles but with a much smaller size.

### Transmission electron microscopy

Adult mice were intracardially perfused with phosphate buffer and a glutaraldehyde/formaldehyde fixative. The thickest segment of the clitoris was isolated and the distal third of the glans penis was microdissected from the internal tissue and flattened on a piece of filter paper for overnight post-fixation at 4 °C. After samples were washed in 0.1M phosphate buffer (pH 7.4), they were cryoprotected overnight in 30% sucrose in 0.1M phosphate buffer. After samples were embedded in Neg-50 Frozen Section Medium, they were frozen and cryosectioned into 100 µm sections and placed into the 0.1M phosphate buffer solution. Medial sections of the clitoris and complete sections of the glans penis were selected for sample preparation.

Samples were osmicated in cacodylate buffer with 1% osmium tetroxide/1.5% potassium ferrocyanide for 1 hour. Sections were washed with ddH2O and stained in a solution of 0.05 M sodium maleate (pH 5.15) and 1% uranyl acetate at 4 °C overnight. After washing with ddH2O, sections were dehydrated with serial ethanol concentrations and propylene oxide. Sections were then infiltrated with 1:1 mix of epoxy resin (LX-112, Ladd Research) and propylene oxide at 4 °C overnight. Sections were embedded in an epoxy resin mix and cured at 60 °C for 48–72 hour. Ultrathin ∼60 nm sections were generated and imaged on a JEOL 1200EX transmission electron microscope at 80 kV accelerating voltage. Images were cropped using Fiji/ImageJ.

### Whole-mount alkaline phosphatase staining of skin and spinal cord

TrkB^+^ and Ret^+^ afferents were sparsely labeled using the *Brn3a^cKOAP^*placental alkaline phosphatase (PLAP) reporter mouse (*44*), as described above, to enable visualization of single nerve terminals in genital and glabrous tissue. Different dissection methods were used for different types of tissue: (1) for female genital tissue, following the clitoris dissection described above, a cut was made along the ventral midline of the urethra to allow spreading of the “wings” of the distal clitoris. (2) For male genital tissue, to permit comprehensive visualization of penile innervation, the superficial layer together with the corpus cavernosum of the glans penis was carefully removed from the internal mesenchyme including the os penis, and separated into two pieces. While not included in our primary analysis, the internal prepuce of the penis was kept for whole-mount alkaline phosphatase staining. To isolate the prostatic urethra, also excluded from our primary analysis, the seminal vesicles and lobes of the prostate were removed, and a cut was made from the distal urethral opening to the opening of the bladder to permit flattening of the tissue. (3) For glabrous skin, the entire ventral surface of the paw was dissected from the underlying tissue for staining. (4) For spinal cord and dorsal column nucleus (DCN), the whole spinal cord was dissected with the DCN attached, and the overlying dura was removed.

The post-fixed and dissected tissue was incubated in PBS at 68°C for 2 hours to inactivate endogenous alkaline phosphatase, then rinsed 3×5 minutes in B3 buffer (0.1M Tris pH 9.5, 0.1M NaCl, 50 mM MgCl2, 0.1% Tween-20) at room temperature. For the PLAP enzymatic reaction, tissue samples were incubated at room temperature overnight in BCIP and NBT in B3 buffer (3.4 µL of each per 1 mL of B3 buffer). Stained tissue was then pinned flat in a dish, fixed in 4% PFA in PBS for 1 hour at room temperature, and serially dehydrated in ethanol (50%, 75%, 100%, 1 hour each, 100% overnight) while covered. Tissue was then cleared and imaged in BABB using a Zeiss AxioZoom stereoscope.

### Quantification and reconstruction of single-neuron morphology

Arborizations belonging to individual neurons were imaged for analysis. The terminal area was measured in ImageJ by drawing a tight polygon around the terminals of a given axon, and the number of corpuscles innervated by each fiber was counted manually. It is possible that the corpuscles of a given afferent occupy a larger volume in the Z dimension than is measured in this manner. Simple corpuscles were deemed to be thin and linear terminals, while complex corpuscles were identified as having wider and bulbous terminals. Reconstruction and filling of representative fibers was completed in ImageJ’s SNT plugin.

### Genital skin injections

Young adult mice were anesthetized with continuous inhalation of 2% isoflurane from a precision vaporizer for the duration of the procedure (5-10 minutes). Injections were done with a beveled borosilicate or quartz glass pipette. The glass pipette was connected to an aspirator tube assembly (Sigma, #A5177-5EA) which was connected to a syringe to control the air pressure.

For penis injections, pressure was applied near the genital region to fully externalize the glans penis from the external prepuce and internal prepuce. A pair of blunt forceps was used to stabilize the penis, while inserting the glass pipette into the distal end of penis. The penetration depth ranges from superficial to 3 mm deep to label the whole glans penis. Pressure was applied to the syringe to eject the liquid inside the glass pipette. After injection, the pipette was kept inside the tissue for 10 s to reduce the leaking. A total of 3 – 4 locations on the penis were injected. For successful injections, the fast green mixed in the liquid was visible inside the tissue for at least 10 min. Rapid dissipation of the fast green dye (within seconds) indicated a failed injection.

For clitoris injections, hairs near the genital protrusion (hairy skin) were removed by Nair treatment and subsequent cleaning using 70% ethanol. A pair of blunt forceps was used to stabilize the protrusion while inserting the glass pipette deep into the middle part of protrusion, ∼1-2 mm. Care was taken to ensure that the needle did not impale the urethra. After injection, the pipette was kept inside the tissue for 30 seconds to minimize leak into the hairy skin region. An injection into the clitoris was considered successful if the fast green mixed in the liquid was visible below but not on the surface of hairy skin.

For injection of mid-line hairy skin, hairs near the genital protrusion in either male or female were removed by Nair treatment and subsequently cleaning using 70% ethanol. A pair of blunt forceps was used to stabilize the hairy skin while inserting the glass pipette into the superficial hairy skin. Following a successful injection, the fast green was immediately visible within the hairy skin region during the injection.

A total volume of 2ul was injected into either the male or female target region, for injection of either CTB or AAV. For CTB injections, 4-8 week old animals were used, and the injected animals were perfused in 3-4 days. For AAV injections, 3-4 week old animals were used, AAV2-retro-hSyn-FlpO (4.95*10^13^, Boston Children’s Hospital Viral Core) was used, and the injected animals were perfused after 3 weeks.

### *In vivo* Multi-Electrode Array (MEA) recordings of L6 DRG neurons

Adult male mice (>6 weeks) were anesthetized with inhaled isoflurane (approximately 2.0%) via a nose cone for the duration of the experiment. Body temperature was maintained at 37°C ± 0.5°C using a custom-made surgical platform equipped with two heating pads mounted on acrylic. An adjustable gap (∼ 1cm wide and ∼7cm long) between the heating pads allowed for access to genital stimulation from below.

After induction of anesthesia, pressure was applied to the abdomen and perineal skin to fully externalize the glans penis. To maintain the externalized state, a minimal amount of Vetbond tissue adhesive was applied to the retracted internal prepuce and the proximal base of glans penis to prevent retraction. Subsequently, the back of the animal was shaved, and a midline incision was made over the lumbar and sacral vertebrae. A custom-made spinal clamp was applied to the L5 vertebrate to secure the spinal column. The paravertebral muscles over the L6 spine were dissected, and the bone covering bilateral L6 DRG was removed with a bone drill. Surgifoam sponges and cotton were used to control bleeding. After cleaning the surface of L6 DRG, the epineurium surrounding the DRG was removed with fine forceps, then saline was added on the top of the DRG.

The platform was then moved to the MEA recording setup. A 32-channel silicon probe (Cambridge NeuroTech, ASSY-37 H6b) was inserted into either the left or right L6 DRG. The signals were amplified and recorded using an Intan Technologies RHD2132 amplifier chip and RHD USB Interface Board. Data acquisition was controlled with open-source software (Intan Technologies Recording Controller version 2.07).

After insertion of the MEA probe into the DRG, brush stimuli using a cotton swab or paintbrush were applied to the glans penis and perineal hairy skin to search for penis-innervating neurons. Neurons that exhibited robust firing in response to mechanical stimulation of the penis but not the adjacent hairy skin were classified as penis-innervating neurons.

For the animals with opsin expression in sensory neurons, the receptive field on the penis was identified, then a fiber optic probe was oriented to deliver light pulses to the penis to optotag sensory neurons expressing opsin. The light pulse was generated by a fiber-coupled LED (M470F4, Thorlabs) with an LED driver (LEDD1B, Thorlabs), and light pulses had a duration of 1ms and a frequency of 10-20Hz. Once entrainment of spiking to light pulses was confirmed, indentation was delivered using a mechanical stimulator (300C-I, Aurora Scientific) with a custom-made indenter using a probe with a ∼200 µm tip diameter. Due to the physical constraint of the setup, only the dorsal side of the penis was accessible for indentation. If the receptive field was on the dorsal side of the penis, the indenter tip was adjusted to the center of receptive field, vertical to the surface. A curved piece of plastic was positioned on the ventral side of the penis to support the indentation. A series of step indentations ranging from 1mN to 75 mN were applied, with each step lasting for 0.5 seconds. A minimum of 20 repeated trials were conducted. Next, a series of sine wave vibrations ramping from 0 to 20mN at varying frequencies (from 10Hz to 120Hz) were applied using the indenter. The vibration stimuli of different frequencies were randomized in order, and each frequency was repeated 5 times. The force and displacement of the indenter were commanded with custom Matlab (version 2019a) scripts controlling a DAQ board (National Instruments, NI USB-6343).

JRCLUST was used to automatically sort action potentials into clusters, which were then manually refined and classified as single or multi units (https://github.com/JaneliaSciComp/JRCLUST).

To calculate the conduction velocity of optotagged neurons, the latency was determined by subtracting the time of each spike from the middle point of each light pulse. The latency for the four optotagged TrkB^+^ Krause afferents were 4.4ms, 4.8ms, 7.8ms, 13.6ms, respectively. The length of axonal projections from L6 DRG to the genitalia in adult mice was estimated to be 5 cm. Thus, conduction velocities of the four optotagged TrkB^+^ neurons are 11.4, 10.4, 6.4, 3.7 m/s respectively.

To determine mechanical thresholds for vibration stimuli across different vibration frequencies, the time of the first spike in response to the vibration in each trial was identified. The corresponding recorded force at that time point was defined as the mechanical threshold for that trial. Thresholds of the trials at each vibration frequency were then averaged.

### *In vivo* calcium imaging of L6 DRG neurons

The surgery procedure for male mice was the same as for the MEA recordings above. After the L6 DRG was exposed, the platform was moved to an upright epifluorescence microscope (Zeiss Axio Examiner) with 10X air objective (Zeiss Epiplan, NA = 0.20) for imaging. The light source was a 470nm LED (M470L5, Thorlabs) with an LED driver (LEDD1B, Thorlabs), and a CMOS camera (CS505MU1, Thorlabs) was triggered at 10 frames per second with 50ms exposure time. All stimuli were synchronized with the camera using a data acquisition board (National Instrument, NI USB-6343).

Indentation on the penis was done in a similar method as described above for the MEA recordings, with a few modifications. The indentation step was increased to 3 seconds to allow for the assessment of adaption properties. The interval between each step was increased to at least 8 seconds to accommodate for the slow decay dynamics of the GCaMP signals. The duration of ramping sinewave vibration was also increased to identify the mechanical thresholds using different vibration frequencies.

For thermal stimulation of the penis, a water reservoir device, inspired by a previously described method (*53*), was used. This device consisted of a reservoir connected to water baths at different temperatures, including room temperature, ice water (∼4°C) and hot water (∼55°C). The penis was submerged in the water reservoir. Then, water of different temperatures was pumped into the reservoir at a controlled speed, and room-temperature water was used during baseline measurements. The water overflowing from the reservoir was directed to a collection chamber located beneath the reservoir. The temperature in the reservoir was monitored using a thermocouple microprobe (IT-1E, Physitemp) and a thermometer (BAT-12, Physitemp).

For clitoris stimulation, the indenter tip was placed vertically on the surface of protrusion where the clitoris was located. To minimize the dorsal-ventral movement of mouse’s lower body during indentation, the mouse tail was held down using a clamp. Similar step indentation and vibration stimuli were used for clitoris as described for the penis. After the mechanical stimulation, the surgery chamber was removed from the microscope, and the anesthetized mouse was placed in a supine position. An incision was created on the protrusion to expose the dorsal nerve of clitoris, and saline was added on the top of the nerve. Subsequently, the mouse was reoriented to the prone position, with the spinal clamp reattached. After locating the same field of view of the DRG, a pair of custom-made bipolar electrodes (silver wire, with ∼1mm distance between the two points) was placed on the dorsal nerve of clitoris, and a train of 1ms pulses (10Hz, less than 1mA) was then delivered.

For the calcium imaging analysis, motion correction and spatial high-pass filtering was conducted using a custom-written ImageJ macro code that employed the ImageJ plugin “moco” and the “Unsharp mask” filter, and then regions of interest (ROIs) were manually selected. Cells exhibiting baseline signal and/or calcium responses were identified and aligned across videos encompassing various stimuli. The intensity measurements generated by ImageJ were then subjected to further analysis using MATLAB. For the calculation of ΔF/F, F was determined using the baseline activity (average fluorescent intensity before each stimulation). The mechanical threshold for each step indentation session was determined based on the first discernible calcium spikes aligned with the step indentation. To determine the mechanical thresholds for vibrations at varying frequencies, the standard deviation of the baseline activity was calculated and the threshold for the response was defined as five times the standard deviation above baseline. Next, the recorded force at that time point was defined and averaged across trials with the same frequency. To differentiate between clitoris-innervating neurons and neurons innervating hairy skin, only neurons that responded to electrical stimulation (five times the standard deviation above baseline) were included for further analysis.

### Spinal transection

Mice were anesthetized with inhaled isoflurane (approximately 2.0%) via nose cone throughout the surgery. The bladder was emptied by applying gentle abdominal pressure. Body temperature was maintained at 37°C ± 0.5°C using a heating pad, and ophthalmic ointment was placed on the eyes. Once reflexes were absent, the back was shaved and sterilized with alternating swabs of ethanol and betadine. A midline incision was made over the thoracic spinal column, and the paravertebral musculature was cut to expose the gap between the T9 and T10 spinal segments. A curved scalpel blade (#10012-00, FST) was inserted between the T9 and T10 spinal segments and moved laterally to ensure thorough transection. Bleeding was controlled with Surgifoam sponges and cotton. The skin of the back was sutured and mice were allowed to recover on a heating pad. Hindlimb paralysis was assessed to confirm success of the spinal transection. Food and DietGel were placed on the floor of the cage, and Carprofen (5mg/kg) was injected subcutaneously every 24 hours. Bladder expression was performed every 2-3 hours on the day of surgery and every 6-8 hours on the following day to prevent urethral blockage by spontaneous plugs. Animals were sacrificed within 3 days of spinal transection.

### Male sexual reflex behaviors

Reflex behaviors were tested at 6 or 24 hours after spinal transection. The awake animal was restrained in a round acrylic chamber in a supine position, with the paralyzed hindlimbs secured on the platform using Scotch tape. Two cameras were positioned from different angles, focusing on the genital region of the mouse. To begin the procedure, pressure was applied to the side of the genital region to fully externalize the glans penis of the mouse. Reflexes might have occured during or right after the externalization due to the force applied in this process, but this effect would end within one minute of externalization.

For evoking vibration-induced sexual reflexes in male mice, two minutes after externalization of the glans penis, a vibrating probe (50 Hz) using a custom-made mechanical stimulator, previously described (*54*), was used. The stimulator consisted of a DC motor connected to an arm (10 mm long) with a round tip (2 mm in diameter) mounted on the end. The motor was driven by a custom-built current supply controlled by a DAQ board (NI USB-6343, National Instrument). The force was calibrated using a custom-made force sensor. The stimulator was attached to an articulating arm that could be moved manually. Before applying vibration stimuli, the stimulator tip was positioned close to the lateral side of glans penis, while a holder attached to an articulating arm was moved to the opposite side to ensure the glans penis remains in place. The vibration stimuli were delivered as a 50 Hz sine wave, with one second of duration followed by a one-second inter-stimulus interval. Each session consisted of 10 such epochs, lasting a total of 20 seconds. The stimuli were programed and delivered using MATLAB. Vibration sessions were spaced two-three minutes apart, and three vibration sessions were conducted for each animal.

For brush-induced sexual reflexes, two minutes after externalization of the glans penis, a cotton swab was used to apply brush stimuli in either the proximal or distal direction. Each session lasted 20 seconds, with intervals longer than two minutes between sessions.

Optogenetic induction of sexual reflexes was performed two minutes after externalization of the glans penis of spinalized animals. The optic fiber (Ø1000 µm, 0.39 NA, M35L02, Thorlabs) connected to a fiber-coupled LED (M470F4, Thorlabs) was directed towards the glans penis. The stimulation sessions lasted 20 seconds, with 10Hz and 2ms pulses used for animals expressing ReaChR, and 20Hz and 1ms pulses used for animals expressing CatCh. The maximum light intensity was 10mW. Animals that showed excessive spontaneous activity before optogenetic stimulation, likely due to a blockage in the bladder, were excluded.

For mechanically induced sexual reflex behaviors, 2–4-month-old animals were used. For the optogenetic stimulation of *TrkB^CreER^* animals, 4-6-month-old animals were used, because externalization of the glans penis was not feasible in younger animals that received tamoxifen treatments at P5. Red light was used to illuminate the setup during optogenetic induced behaviors, while room light was used for mechanical stimulation measurements. To quantify the sexual reflexes, the start and end of each reflex behavior was manually labeled using BORIS software (*55*). The behavioral measurements were then exported, analyzed, and plotted in MATLAB.

### Mating behaviors

At least two weeks prior to testing, mice were transferred to a reverse 12h light/dark cycle room. All animals used for mating behaviors were 2-4 months old. They were from a mixed background but backcrossed to C57Bl/6 mice for two generations.

The mating behavioral analysis was similar to that of a previous study (*56*). The male mouse was singly housed for at least two weeks before testing and was socialized with a female mouse overnight at least one week prior to the test. Two days before the test, female mice were treated with 18-35 μg of 17β-estradiol benzoate (dissolved in sterile sesame oil, final volume: 50-100 μl, subcutaneous injection), and treated again with 50-100 μg progesterone (dissolved in sterile sesame oil, final volume 50-100 μl) five hours before the test. During the mating test, a female was placed into the male’s home cage, and the cage cover was replaced by a custom-made wall (6 inches tall) to prevent animals from leaving their cage. Mating behaviors were visualized in the male’s home cage using USB cameras, which captured videos at 30fps, that were placed above the cage. The behavior room was dark, illuminated only with infrared lamps.

When testing the mating performance of the males, a wild-type female mouse which was unreceptive 10 minutes after the start of the session would be replaced with another female, so that only receptive females were used for testing males. Mating behavior was monitored over video for 75 minutes. The males, starting as sexually inexperienced animals, underwent testing for at least three rounds, with each round occurring at least one week apart.

When testing the mating performance of females, only experienced males were used. The same group of female controls and mutants were hormone-primed (as described above) in the first two rounds of the mating test, while in the third round they were in a natural estrus cycle. The first two rounds were conducted two weeks apart, and the third trial was conducted four weeks after the second trial to allow for the recovery of estrus cycle. The estrus cycle was monitored using the vaginal cytology method, following previously established protocols (*57, 58*). Vaginal smears were collected, stained with crystal violet, and examined to determine the estrus stage. Only female mice in proestrus or estrus were used for mating behaviors within three hours of vaginal lavage. The female was separated from the male 10 minutes after the start of intromission to avoid pregnancy. If no intromission occurred, the mating session was stopped at 30 minutes.

The videos of mating behaviors were manually scored using BORIS software (*55*) by an experimenter who was blinded to the genotype of the animals. For male mating behavioral analysis, the following behaviors were scored: sniffing, mounting, intromission, and ejaculation. Mounting refers to the males’ rapid pursuit of females, followed by grasping their rear, and often followed by short probing without gaining access to the female genitalia. Intromission represents the long rhythmic thrusting, indicating the male’s successful penetration of the vagina. The mounting and intromission periods do not overlap with the scoring methods used; mounting is considered the pursuit and adjustment period prior to the intromission period. The intromission bouts lasting less than 2 seconds were excluded from the analysis due to the lack of rhythmicity of movements and the possibility that penetration did not fully occur. In addition, combative behaviors and darting behaviors of female mice were scored during the female mating behavior assessment trials. Combative behavior refers to the instances where the female confronts or fights with the male, while the dart behavior occurs when the female mouse attempts to quickly escape from the mounting of the male mice.

---

**Fig. S1.**
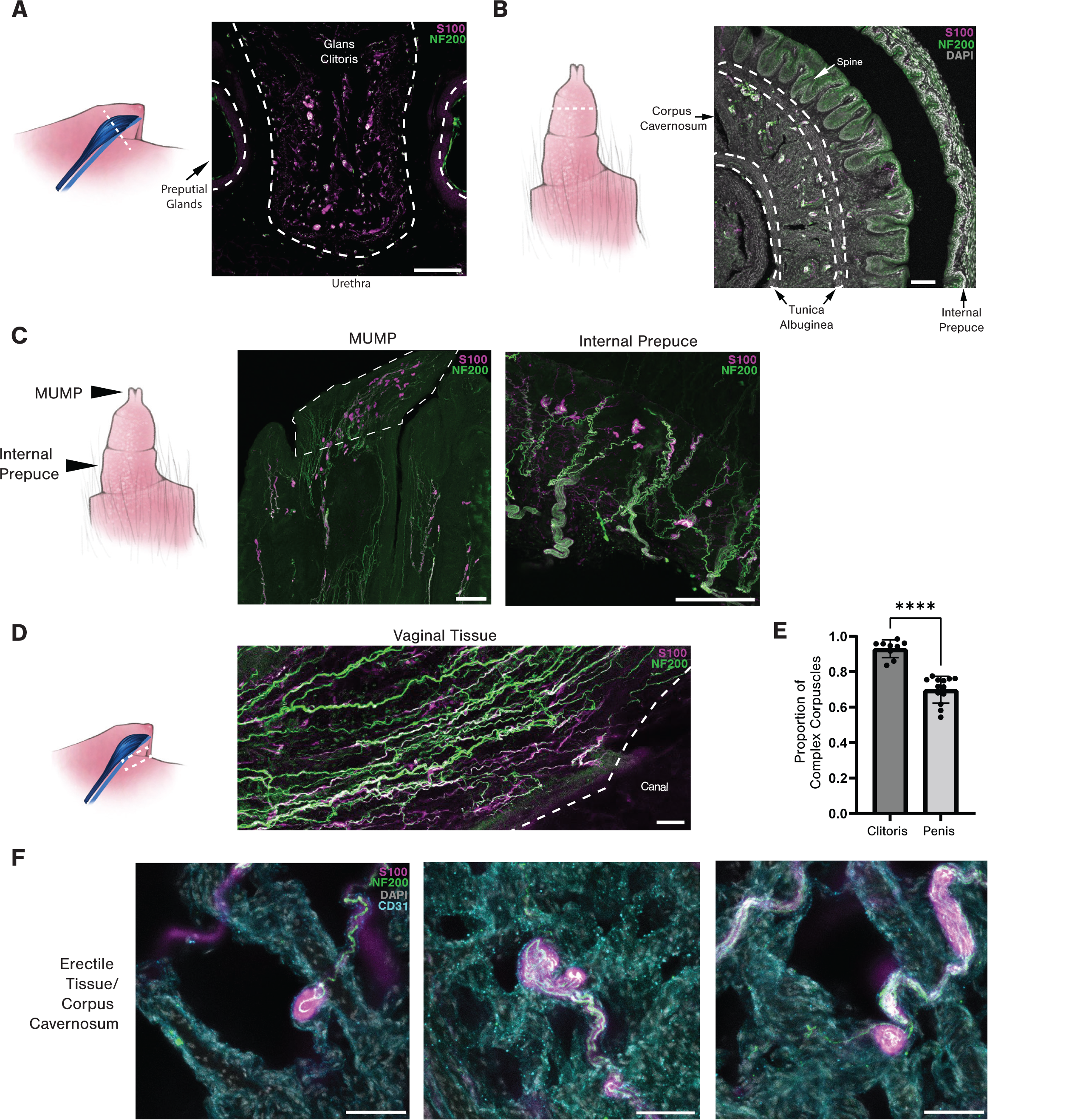
Distribution of Krause corpuscles. (**A**) Immunostaining of a coronal section through the clitoris and surrounding tissue (dashed line) for S100 and NF200. The clitoris, urethra, and preputial glands are annotated. (**B**) Immunostaining of a coronal section through the penis (dashed line) as in (A). The internal prepuce, penile spines of the glans penis, corpus cavernosum, and surrounding tunica albuginea are annotated. (**C)** Corpuscles within the male urogenital mating protuberance (MUMP) and internal prepuce are visualized as in (A). (**D**) Immunostaining of a section of vaginal tissue, stained as in (A), isolated from the dashed area, shows an absence of Krause corpuscles. (**E**) Quantification of the percent of complex Krause corpuscles among the total corpuscles observed in a section of genital tissue (n = 9 sections from 4 females and 13 sections from 4 males). ****p < 0.0001, unpaired t-test. (**F**) Immunostained sections showing Krause corpuscles within the corpus cavernosum of erect penis tissue for S100, NF200, and CD31. Scale bar is 100 µm in A-D and 50 µm in F.

**Fig. S2.**
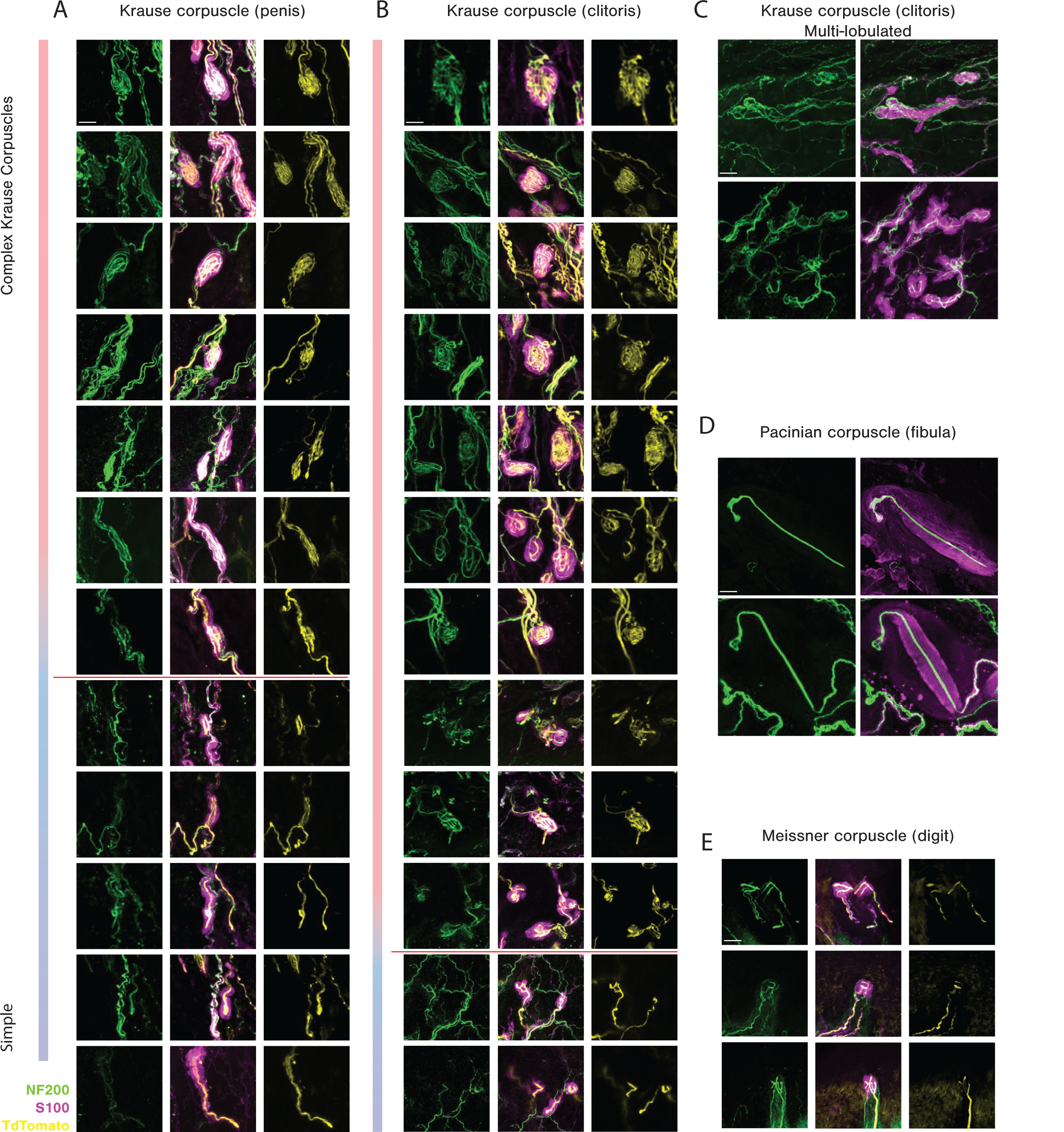
The diversity of Krause corpuscle morphologies contrasts with Meissner and Pacinian corpuscles located in other body regions. The range of Krause corpuscle morphologies found in the penis (**A**) and clitoris (**B**), from complex (top) to simple (bottom). The red line depicts the distinction between complex (above the line) and simple (below the line) Krause corpuscles for quantification in fig. S1E. (**C**) Examples of multi-lobulated Krause corpuscles observed in the clitoris. (**D**) Examples of Pacinian corpuscles located in the periosteum of the fibula. (**E**) Examples of Meissner corpuscles located in dermal papilla of glabrous skin of a forepaw digit tip. The scale bar is 20 µm, and all images are of the same scale. Images in A, B and E are stained for NF200 (green), S100 (magenta), and tdTomato for TrkB axons (yellow) in *TrkB^CreER^; Avil^FlpO^; R26^FSF-LSL-Tdtamato^* mice. Images in C and D are stained for NF200 (green) and S100 (magenta).

**Fig. S3.**
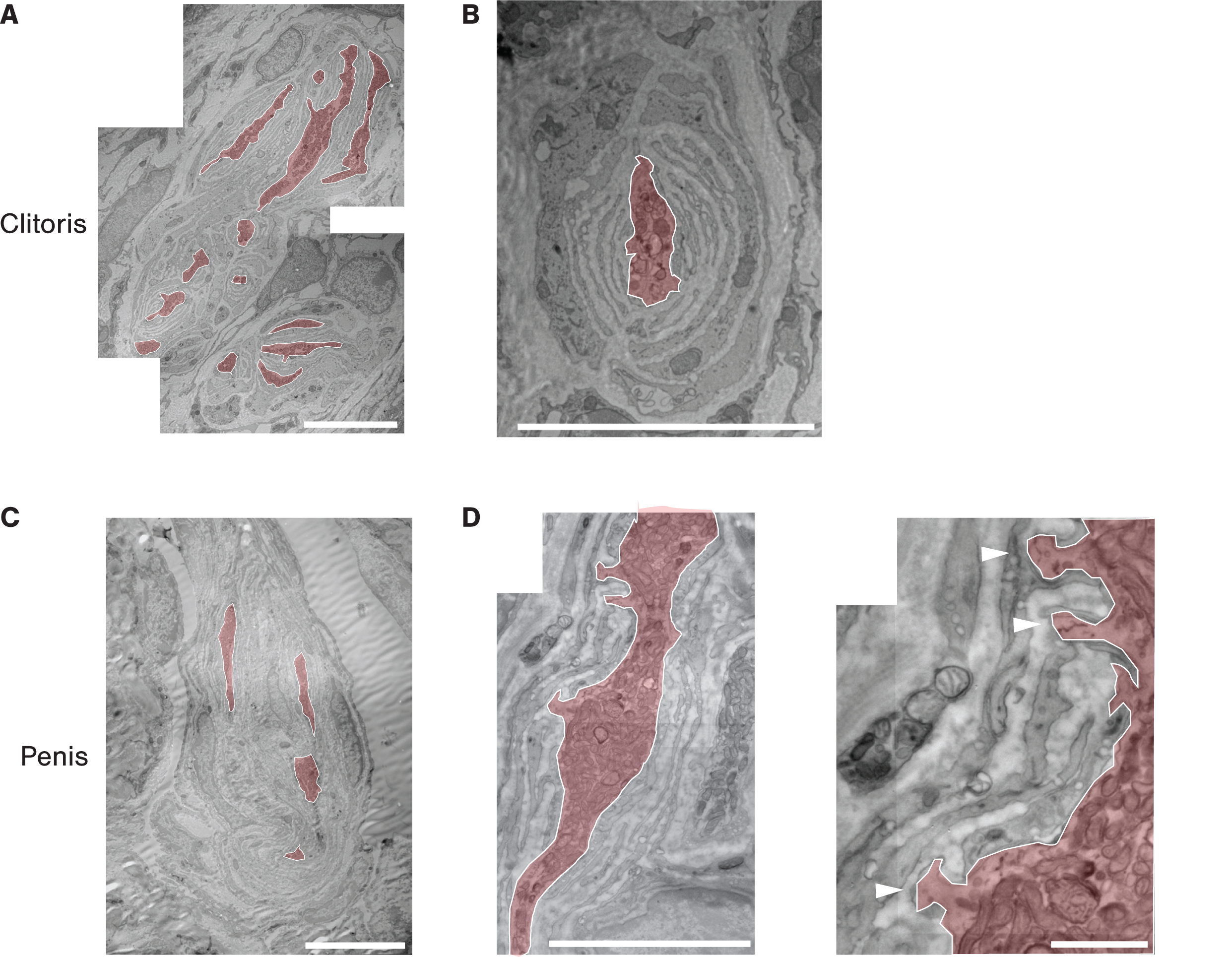
Electron microscopy visualization of Krause corpuscles. (**A**) Representative transmission electron micrograph of a multi-lobulated complex Krause corpuscle within the clitoris. (**B**) Example electron micrograph of a simple Krause corpuscle with a single lamellated axon profile within the clitoris. (**C**) Example electron micrograph of a complex Krause corpuscle within the penis. (**D**) Example electron micrograph of a Krause corpuscle (left) magnified to visualize axonal protrusions (right, white arrowheads). All axon profiles were manually outlined and filled (light red). Scale bar in A-C (left) is 10 µm, D (left) is 5um, D (right) is 1 µm.

**Fig. S4.**
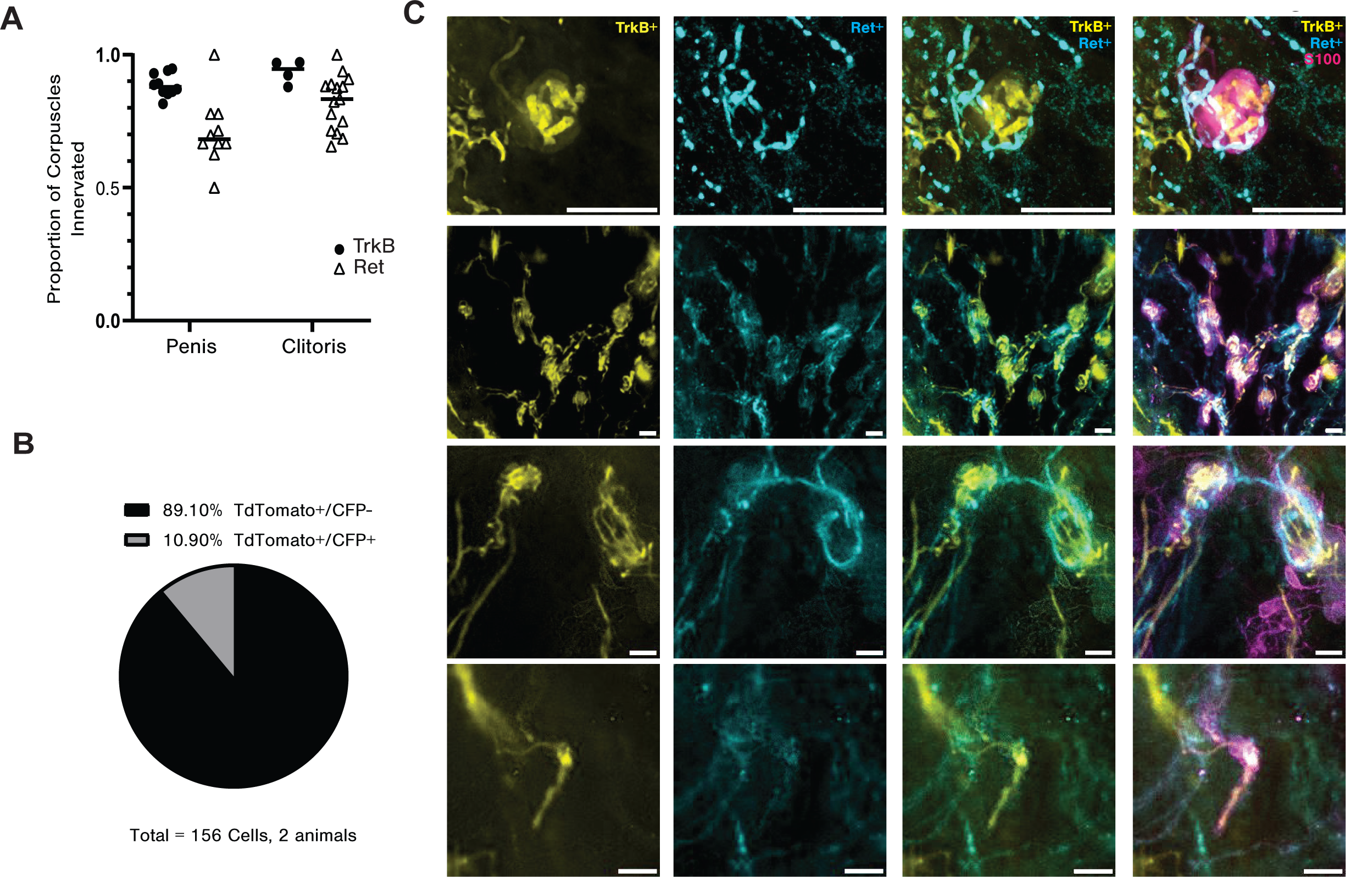
Dual innervation of Krause corpuscles by TrkB^+^ and Ret^+^ DRG sensory neurons. (**A**) Quantification of the fraction of corpuscles innervated by NF200^+^ axons and labeled using *TrkB^CreER^; Avil^FlpO^; R26^FSF-LSL-Tdtamato^*or *Ret^CFP^* mice, among all S100^+^ corpuscles in a given section (TrkB: 10 sections from three males [89.4 ± 4.2%] and four sections from two females [93.5 ± 4.4%]; Ret: 10 sections from two males [70.9 ± 12.3%] and 15 sections from three females [80.0 ± 10.2%]). (**B**) Quantification of the overlap of TdTomato and CFP in DRG soma of *TrkB^CreER^*; Ai14; *Ret^CFP^* mice. (**C**) Dual labeling of Ret^+^ and TrkB^+^ Krause corpuscle afferents showing dually-innervated corpuscles and a simple corpuscle innervated only by a TrkB^+^ fiber (bottom) stained for tdTomato (TrkB^+^), GFP (Ret^+^), and S100. Scale bar is 10 µm in the panels of the first row and 50 µm in all other panels.

**Fig. S5.**
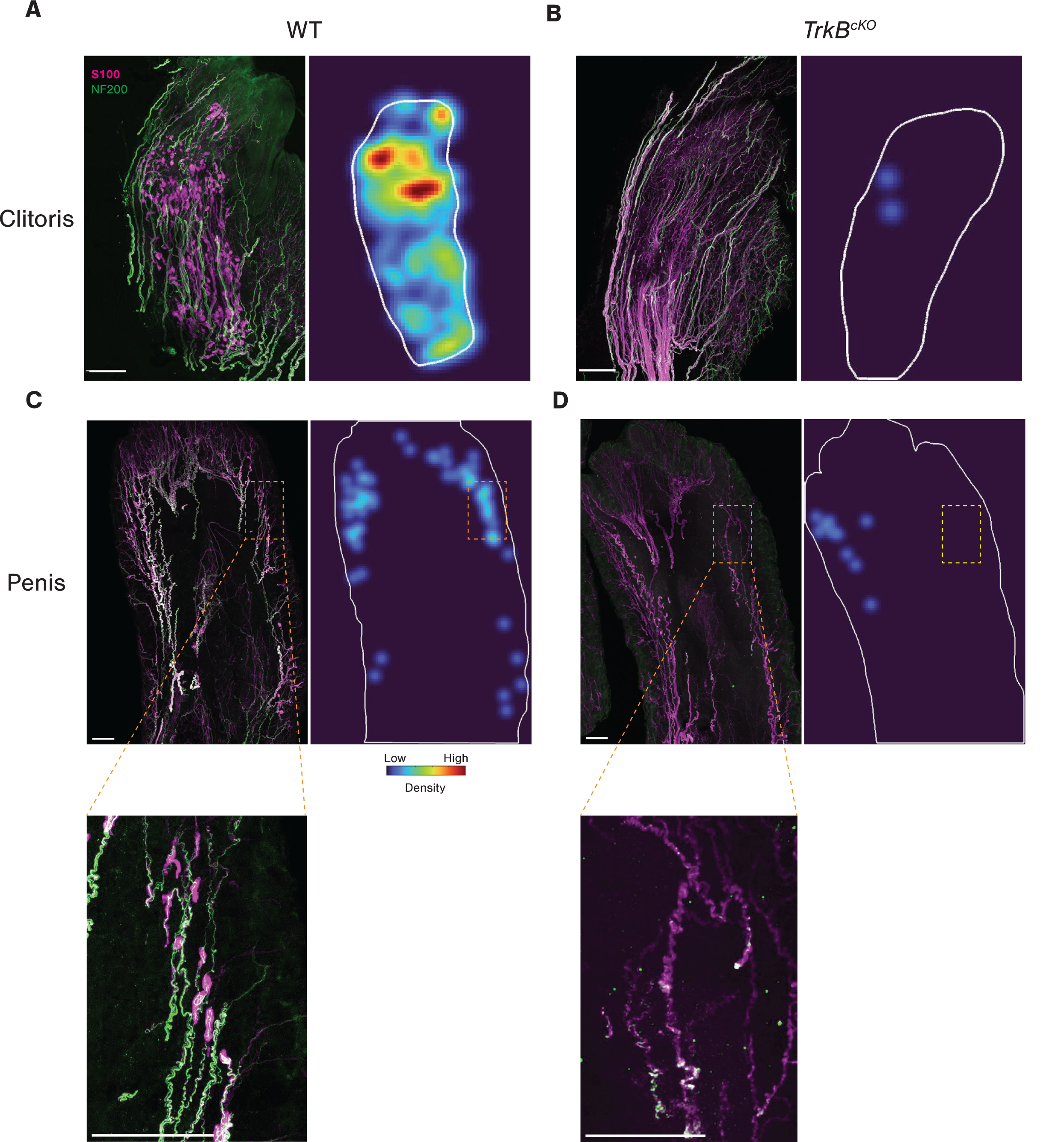
*TrkB^cKO^* animals lack Krause corpuscles in both the clitoris and penis. Representative images of 200 µm-thick sections of the clitoris (**A and B)** and glans penis (**C and D**) of wild type (A and C) and *TrkB^cKO^*(B and D) animals stained with NF200 and S100. For images of the glans penis, the regions indicated by rectangles are magnified below. To the right of each image is the corresponding density heatmap of Krause corpuscle distributions. The border of the section is outlined in white. The same color scale, normalized by the densest area of the clitoris, is used for all four heatmaps. Scale bar = 200 µm.

**Fig. S6.**
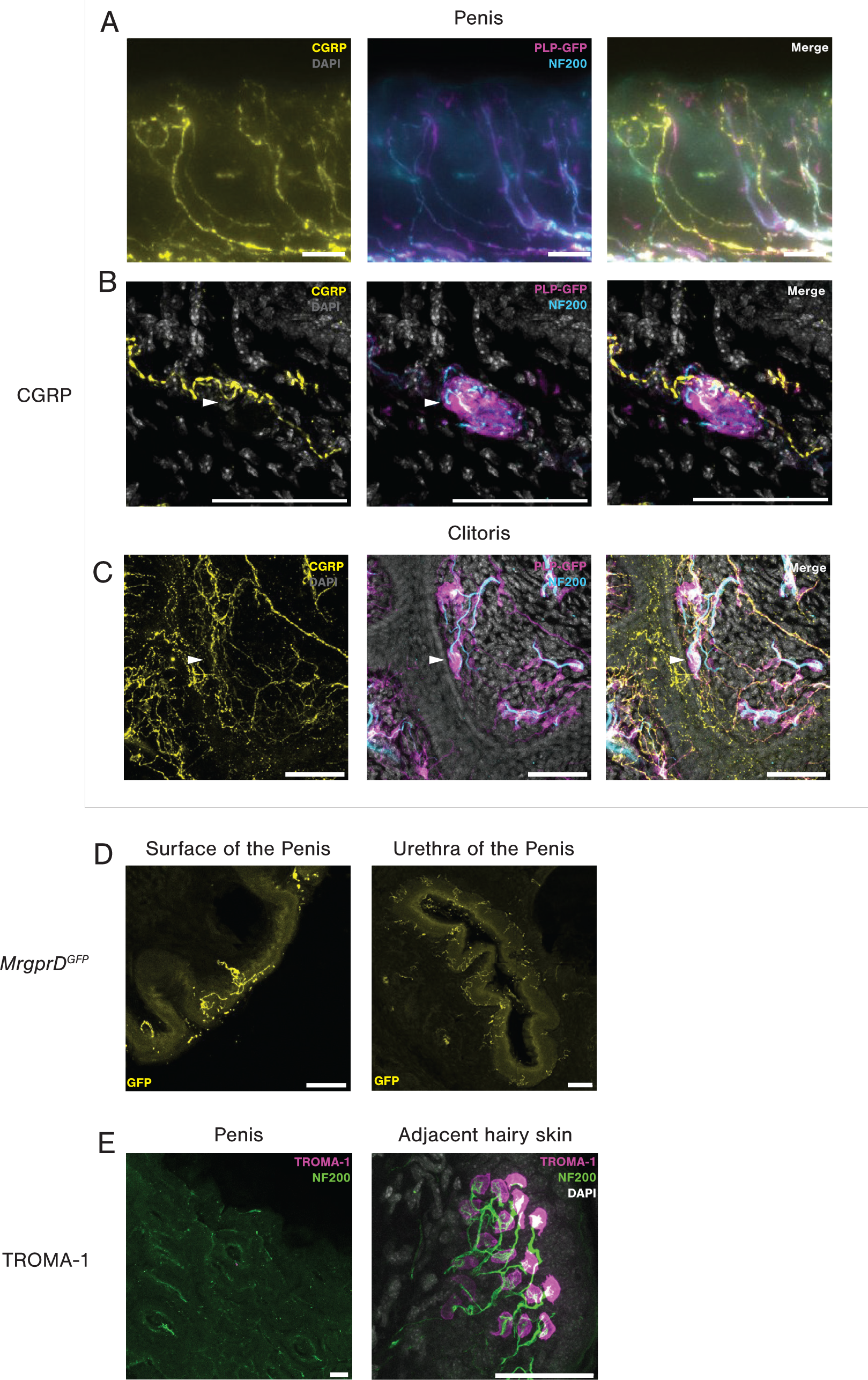
CGRP^+^ and MrgprD^+^ fibers form free nerve endings in female and male genital tissue. (**A** to **C**) Representative images of immunostaining glans penis sections (200 µm thick) (superficial tissue in A, deeper tissue in B) and clitoris sections (200 µm thick) (C) for CGRP, NF200, and GFP to label Schwann cells in *PLP^EGFP^* animals. The white arrowheads in B and C indicate Krause corpuscles that have CGRP^+^ fiber passing through or terminating within them. (**D**) GFP immunostaining of coronal sections of the glans penis from *MrgprD^GFP^* animals. Left: superficial free nerve endings; right: nerve terminals near the urethra of the penis. (**E**) Representative images of co-staining of TROMA-1, a marker of Merkel cells, and NF200. Left: a section of glans penis; right: a section of adjacent hairy skin. Scale bar in all images is 50 µm.

**Fig. S7.**
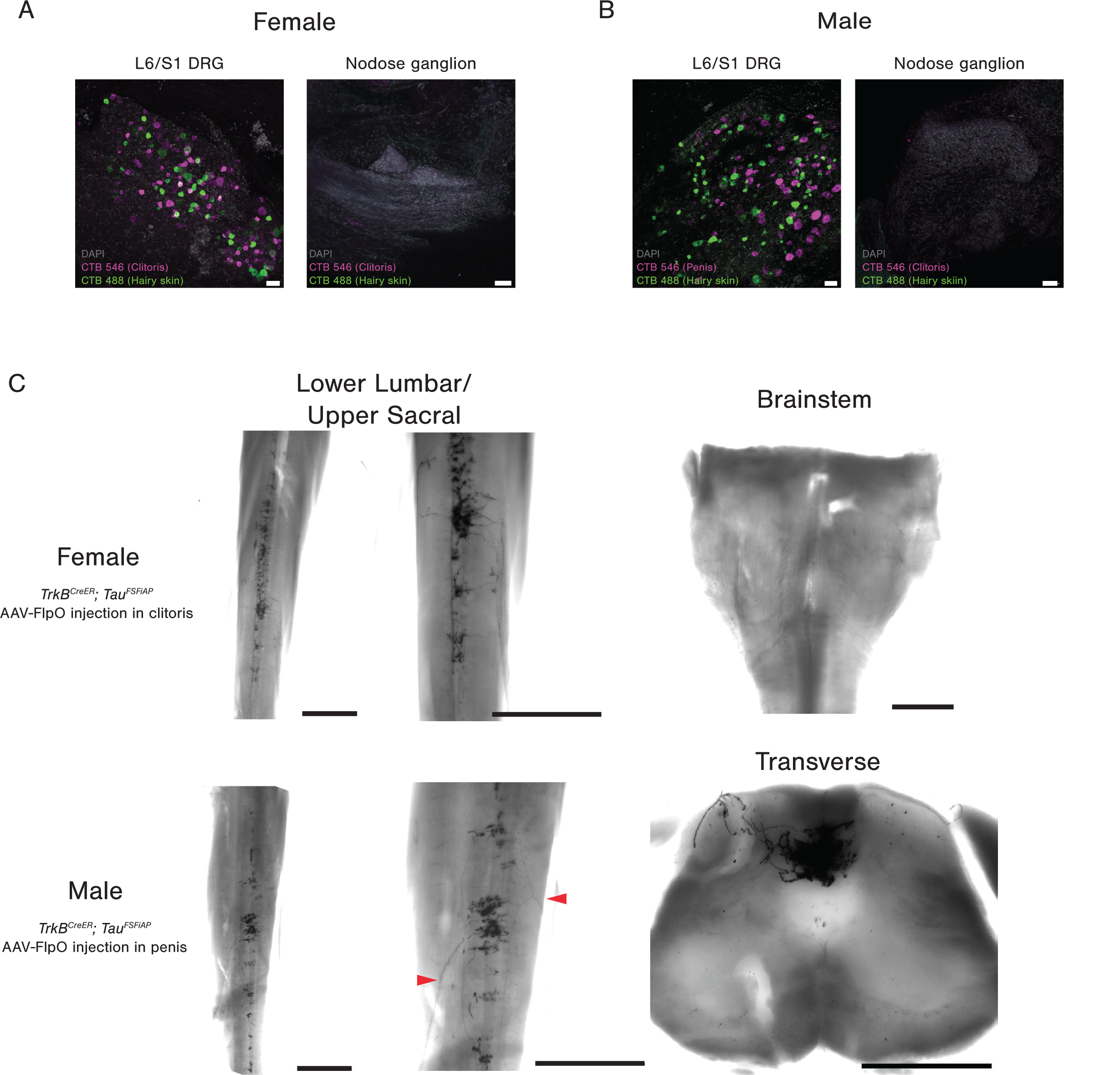
The central terminals of Krause corpuscle afferents. (**A** and **B**) Representative images of whole-mount L6 or S1 DRGs or nodose ganglion after injection of CTB tracers conjugated to different fluorophores into genital tissues (CTB-546) and adjacent mid-line hairy skin (CTB-488). Scale bar is 50 µm. (**C**) Whole-mount AP staining of the spinal cord of *TrkB^CreER^; Tau^FSFiAP^* animals with AAV2-retro-hSyn-FlpO injected into the clitoris (upper panels) or penis (lower panels). The middle panels show magnified images of those shown on the left. Red arrowheads point to the individual axons entering the spinal cord. The images on the right show whole mount AP staining of the dorsal column nucleus (top) and a transverse section of the S1 spinal cord (bottom). Scale bar in (C) is 1 mm for whole mount images and 500 µm for transverse sections of AP staining.

**Fig. S8.**
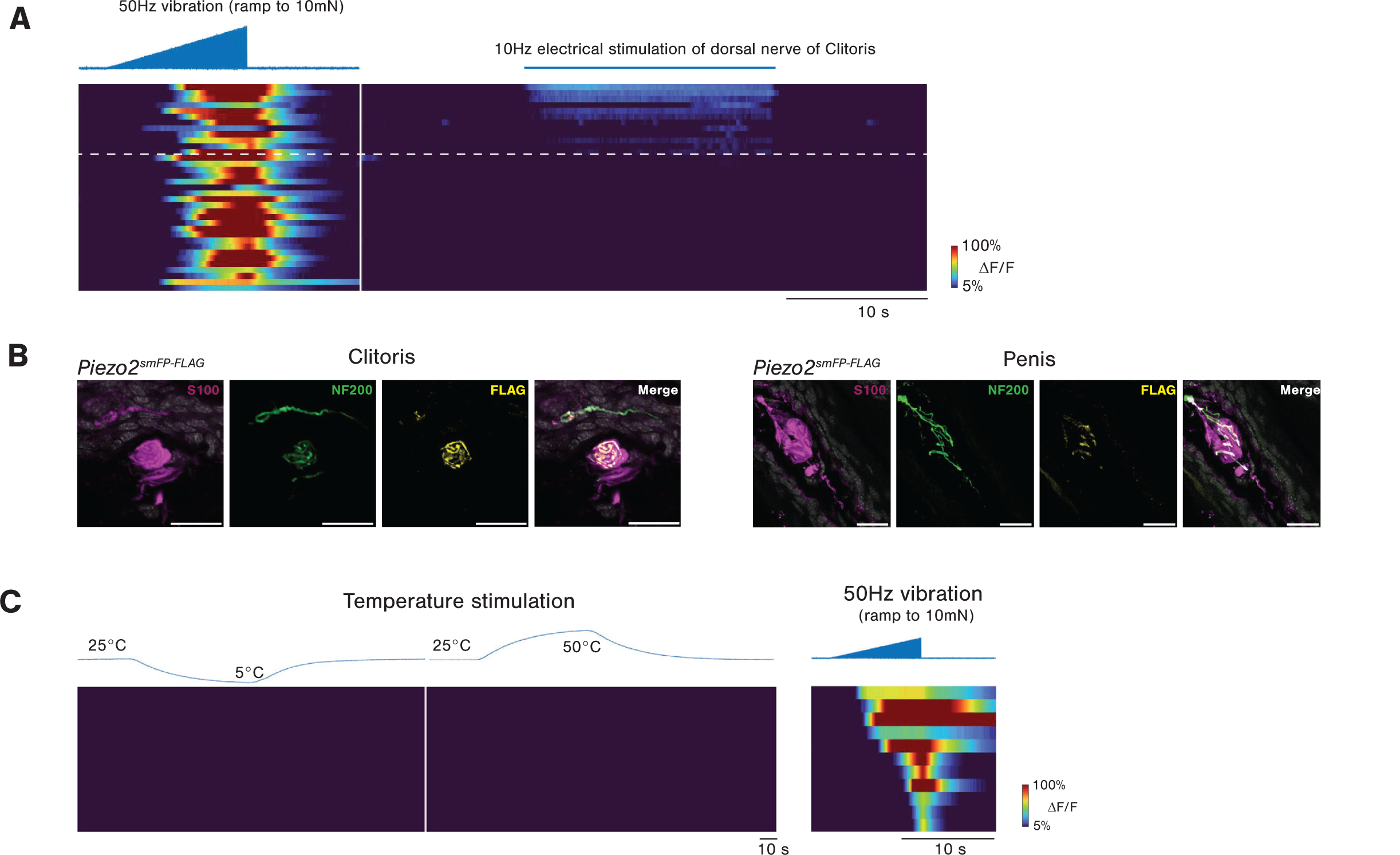
Krause corpuscle afferents are mechanosensitive neurons tuned to dynamic, light touch and mechanical vibration. (**A**) Calcium signals of TrkB^+^ neurons in L6 DRG in response to ramping vibration stimuli applied to the protrusion where the clitoris is located (left) and to electrical stimulation of the dorsal nerve of the clitoris (right). Neurons responding to the electrical stimulation were identified as clitoris-innervating neurons (above the dashed line). The neurons below the dashed line are presumed to innervate the adjacent hairy skin. The neurons shown in (A) are from three female mice. (**B**) Representative images of Piezo2 expression in the axons innervating Krause corpuscles from *Piezo2^smFP-FLAG^* animals. Sections were immunostained for S100, NF200, and FLAG. Scale bar: 50 µm. (**C**) Left: thermal responses of TrkB^+^ afferents that innervate the penis. The top trace shows the measured temperature of the water bath in which the penis was submerged. Right: the response of the same group of neurons to vibration applied to the glans penis.

**Fig. S9.**
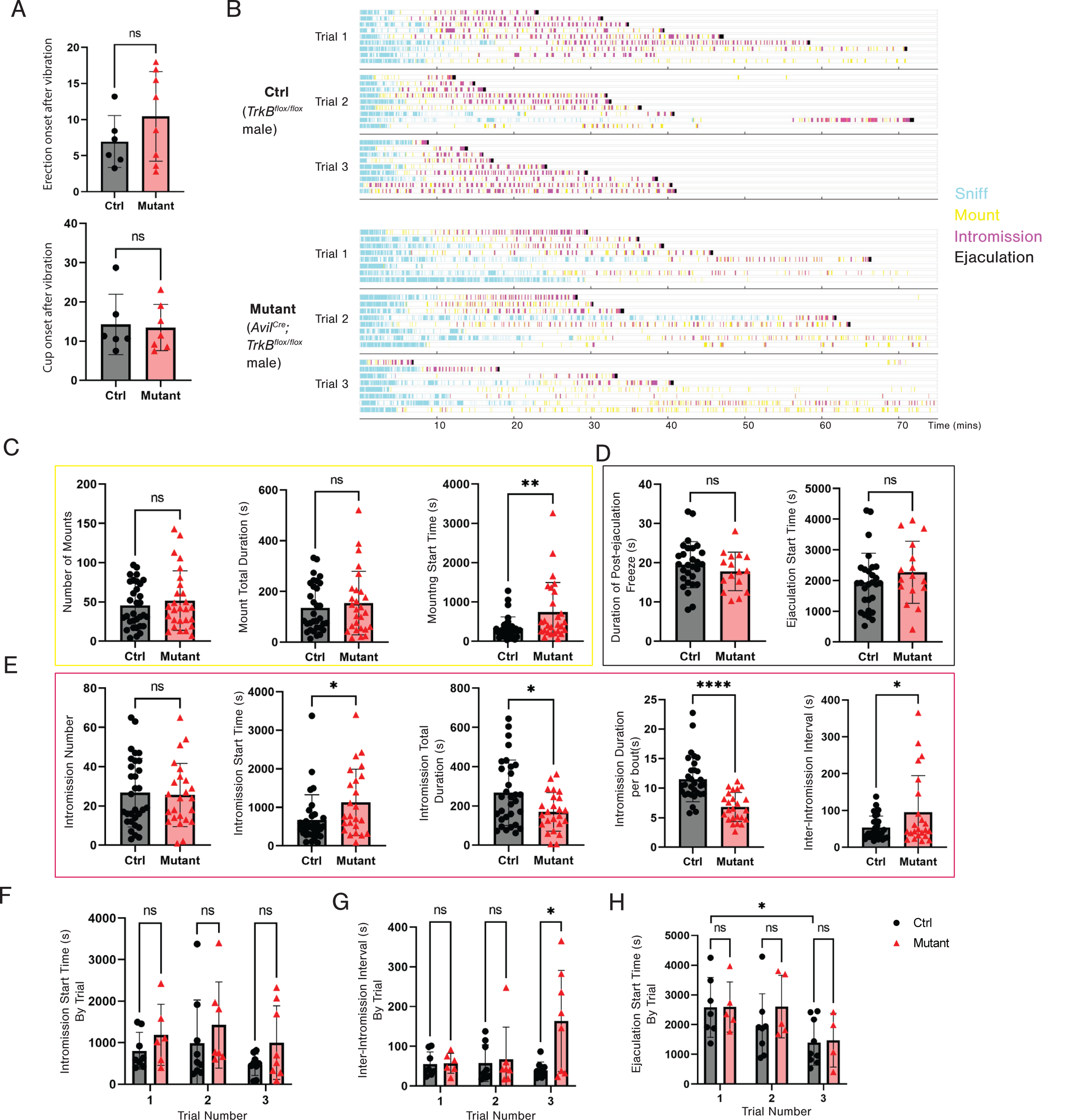
*TrkB^cKO^* males display deficits in mating behaviors. (**A**) Genital reflexes following 50 Hz vibration applied to the penis of control (ctrl, n=6) and *TrkB^cKO^* males (n=7) that have undergone spinal transection. Unpaired t-test, p>0.05. (**B**) Ethograms of mating behavior of *TrkB^cKO^* (n=8) and littermate control (n=9) animals within a 75-minute testing period across three trials, with trials separated by at least one week. The sniffing bouts shown are limited to the male’s sniffing behavior before the first mount. Mounting refers to shallow probing and postural maneuvering without gaining access to the female genitalia, while intromission represents periods of time during which long rhythmic thrusts are observed. (**C**) Quantification of mounting behaviors across three trials. Unpaired t-test, ** p<0.01. (**D**) Quantification of ejaculation behaviors, collected across three trials. Only animals that achieved ejaculation are included. (**E**) Quantification of intromission behaviors across three trials. Unpaired t-test, * p<0.05, **** p<0.0001 (for intromission start time, * p = 0.029; for intromission total duration, * p = 0.014; for inter-intromission interval, * p = 0.028). (**F-G**) Comparison of intromission start time (F), inter-intromission interval, and ejaculation start time within each trial. Unpaired t-test, * p<0.05.

**Fig. S10.**
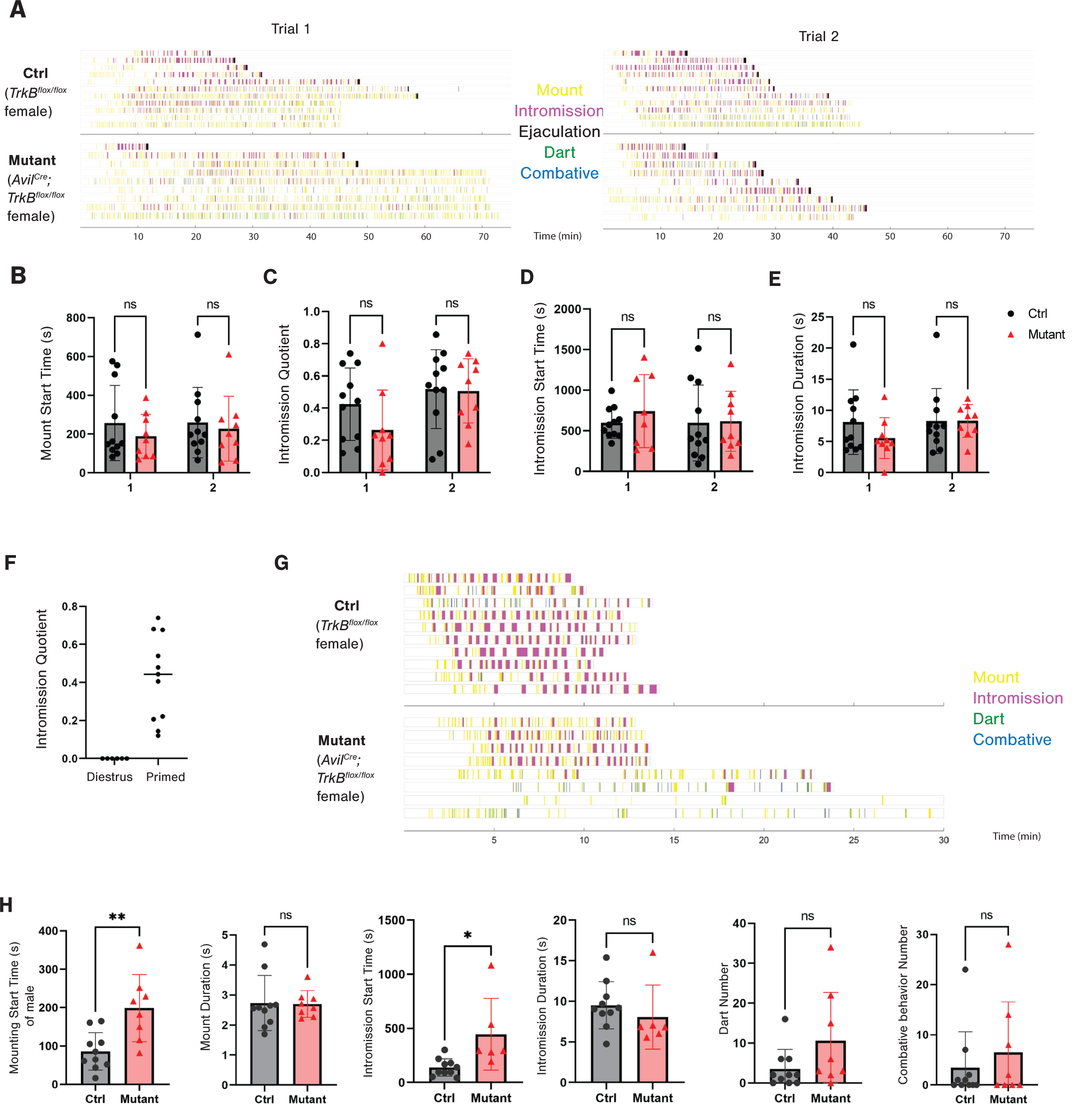
*TrkB^cKO^* females display deficits in mating behaviors. (**A**) Ethograms of mating behaviors of hormone-primed *TrkB^cKO^*(n=9) and littermate control (n=11) females within a 75 minute testing period across two trials, with each trial two weeks apart. (**B-E**) Quantifications of mount start time (B), intromission quotient (total intromission time/(total intromission time + total mount time)) (C), intromission start time (D), and average intromission duration per bout (E), within each trial. Unpaired t-test, p>0.05. (**F**) Comparison of intromission quotient of wild-type females in diestrus (without hormone priming) and with hormone priming. (**G**) Ethogram of the mating behavior of naturally-cycled *TrkB^cKO^* (n=8) and littermate control (n=10) females. Females were separated from males 10 minutes after the start of intromission to avoid pregnancy. If no intromission occurred, the session was stopped at 30 minutes. (**H**) Quantifications of mating behavior of naturally-cycled *TrkB^cKO^* (n=8) and littermate control (n=10) females. Unpaired t-test, * p<0.05, ** p<0.01.

